# OxyBLI: A Genetics-Based Approach for *In Vivo* Real-Time Visualization of Tissue Oxygenation Dynamics

**DOI:** 10.64898/2026.06.24.734154

**Authors:** Satoshi Iwano, Jungo Kato, Tomoko Toramaru, Hiroshi Hama, Mayu Sugiyama, Reiko Takahashi, Megumu Takahashi, Hiroyuki Hioki, Toshiaki Nakashiba, Atsushi Miyawaki

## Abstract

Accurate measurement of cellular oxygen levels is essential for understanding the balance between oxygen demand and supply in tissues. However, conventional methods only yield compromised results. We harnessed the oxygen dependence of bioluminescence to develop OxyBLI—a noninvasive optical method that directly monitors oxygen levels in specific cell populations of intact experimental animals. We characterized OxyBLI signals across various critical situations associated with common interventions. Hypoxic breathing and subsequent systemic tissue hypoxia caused blood to be redistributed in a way that prioritized brain oxygenation. In contrast, hyperoxic breathing sharply increased tissue oxygenation, which promptly returned to the target level owing to a vasoconstrictor response. These findings are expected to help resolve the long-standing clinical issue regarding the risks and benefits of administering supplemental oxygen to acutely ill patients. Our multifaceted approach, which presents multiple challenges to individual animals over time, will advance our understanding of the delicate interaction between hypoxia and hyperoxia.

## Introduction

Molecular oxygen (O_2_) concentration in the air is currently 20.8%, but it was extremely low on very early Earth. O_2_ accumulation in the atmosphere began around 2.4 billion years ago (the Oxygen Catastrophe or Holocaust)^1,2^. Some organisms evolved cleverly by developing mechanisms to detoxify and exploit O_2_, enabling them to survive in the new aerobic environment. Conversely, organisms unable to adapt to O_2_ were relegated to anaerobic niches. Atmospheric O_2_ levels fluctuated over time, reaching a high of 35% during the Carboniferous period, roughly 300 million years ago^3^. This elevated O_2_ level is thought to have contributed to the size increase of animals at that time. Thus, until the end of the 18^th^ century, when humankind discovered O_2_, most life in the biosphere had never experienced hyperoxic environments exceeding 35%. Today, we have the opportunity to be exposed to artificially produced pure O_2_ gas. Supplemental O_2_ administration is now widely used in emergency and intensive care medicine, and demand for O_2_ therapy markedly increased during the Coronavirus pandemic.

Currently, O_2_ is essential for the cellular respiration of aerobic organisms on Earth, including mammals. Low levels of O_2_, or hypoxia, result in severe complications, including organ dysfunction, brain damage, and cardiac arrest. Numerous studies have revealed that hypoxia is one of the most powerful inducers of gene expression, metabolic changes, and regenerative processes^4,5^. Blood oxygenated by the respiratory system is conveyed throughout the body by the circulatory system for tissue oxygenation (Fig. 1). Hypoxemia (low levels of O_2_ in the blood) is easily monitored by pulse oximetry, a noninvasive method for measuring O_2_ saturation through the skin (SpO_2_). By contrast, the pathophysiology of hyperoxia, a state in which the O_2_ supply to tissues is excessive, has remained much less explored. Although hyperoxic breathing or O_2_ inhalation to promote tissue oxygenation is a common clinical intervention for critically ill patients, its safety and effectiveness have been debated^6–8^. As SpO_2_ cannot detect high levels of blood O_2_ (hyperoxemia), the amount of O_2_ administered is extremely difficult to titrate. Consequently, many patients in crisis are likely exposed to episodes of hyperoxemia. Then, the question is to what extent hyperoxemia causes hyperoxia in tissues. Tissue hyperoxia may induce tissue damage by producing reactive O_2_ species (ROS) and causing oxidative stress. In fact, liberal O_2_ administration was found to be associated with increased mortality in acutely ill adults^9^. However, this intervention is assumed to be countered immediately by the vasoconstrictor responses of small arterioles to O_2_, thereby preventing an undesirable increase in tissue oxygenation. Nevertheless, O_2_ inhalation remains the most effective way to improve tissue hypoxia. Many uncertainties in clinical studies stem from the lack of a method to directly measure tissue O_2_ levels.

**Figure 1.**
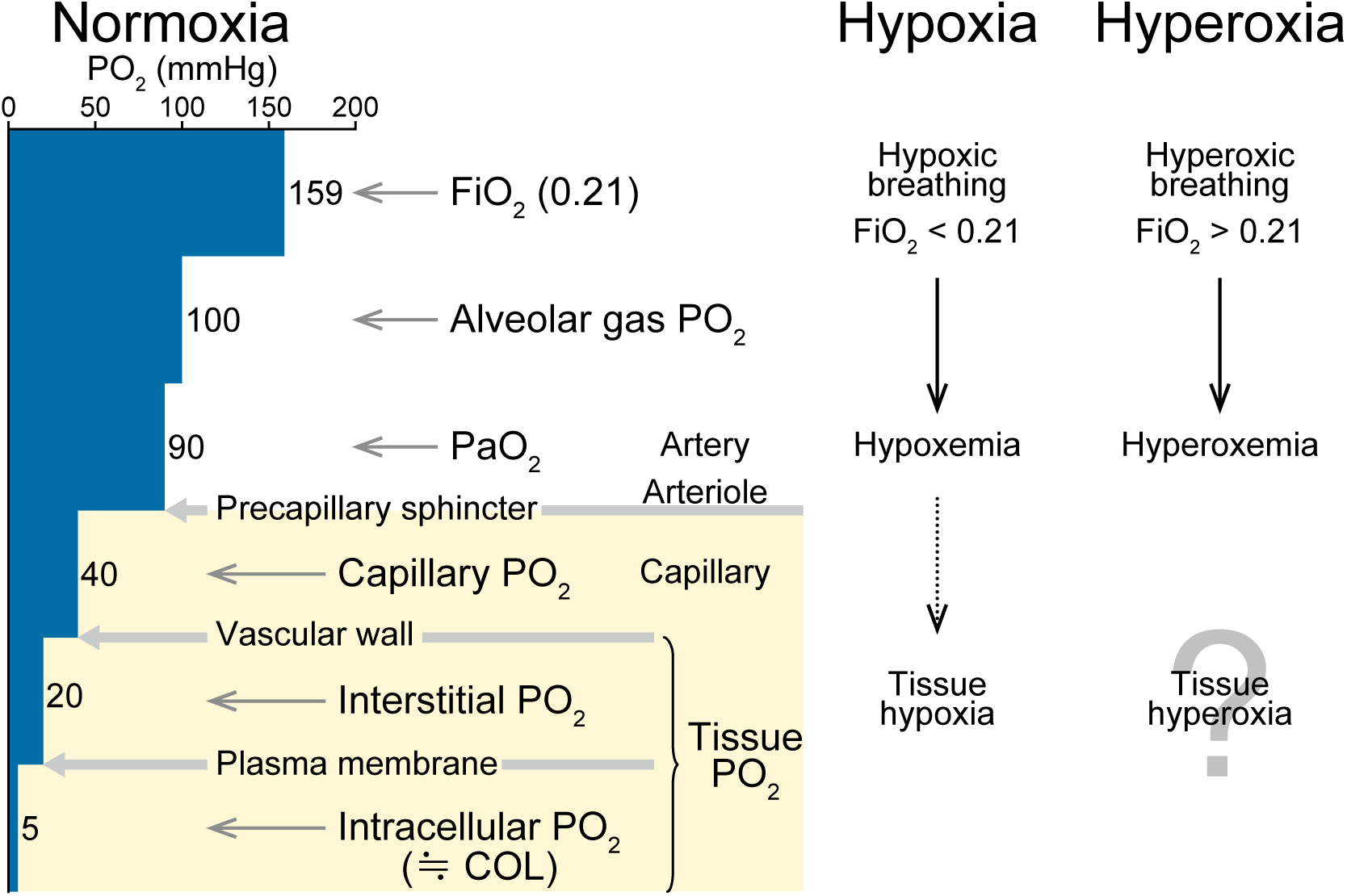
O_2_ distribution in the respiro-circulatory system and tissues of the body in normoxic, hypoxic, and hyperoxic environments. There is a longitudinal gradient of PO_2_ in the arterial network, likely due to O_2_ consumption in the arterial wall. Additionally, precapillary arterioles regulate O_2_ transport from arterial blood into tissues. The figure illustrates the stepwise decline in PO_2_ along with the average values (blue bars). While PO_2_ in arterial blood (PaO_2_) is around 90 mmHg, PO_2_ in capillary blood is estimated to be roughly 40 mmHg. While PaO_2_ is directly measured by blood gas analysis in clinical practice, measuring capillary PO_2_ is extremely challenging. In this study, PaO_2_ was continuously monitored by inserting a fiber-optic O_2_ sensor into the femoral artery of anesthetized rats. Tissue O_2_ is further categorized into interstitial and intracellular O_2_. Conventional tissue PO_2_ recordings reflect the average PO_2_ in interstitial fluids. In contrast, COLs (cellular O_2_ levels) measured in this study reflect the average PO_2_ inside cells. In principle, hypoxia and hyperoxia refer to states in which O_2_ supply to the tissues is insufficient and excessive, respectively. However, these two states are not simply opposites of each other. The most common type of hypoxia is hypoxic hypoxia or generalized hypoxia, which results from the defective oxygenation of blood in the lungs. For example, a low O_2_ environment (low FiO_2_) results in hypoxemia (low PaO_2_) and, consequently, tissue hypoxia. In contrast, hyperoxia scenarios are complex. Although a high O_2_ environment (high FiO_2_) certainly results in hyperoxemia (high PaO_2_), the consequence may not necessarily be tissue hyperoxia (indicated by a question mark). Due to this uncertainty, hyperoxia usually implies an environment with a high O_2_ concentration. In contrast, hypoxia is mainly discussed on the tissue level. In normoxic conditions, O_2_ is carried primarily by hemoglobin (Hb), and the amount of dissolved O_2_ is relatively small. In hyperoxic conditions, however, the amount of dissolved O_2_ becomes significant, resulting in an elevated PaO_2_. Under such circumstances, the difference between PaO_2_ and tissue PO_2_ becomes more critical than usual for the rate of O_2_ delivery to tissues. For this reason, this figure spotlights PaO_2_ rather than Hb–O_2_ saturation. The compartment protected from an excess O_2_ is shaded in ocher. Downward solid arrows indicate almost fixed consequences. A downward dotted arrow signifies complex regulations, including compensatory adjustments against acute systemic hypoxia, which involve the preferential redistribution of blood flow to the brain. Abbreviations: FiO_2_: fraction of inspired O_2_; PO_2_: partial pressure of O_2_.

To analyze such dynamic responses to inhaled O_2_, it is necessary to image tissue O_2_ levels in real time, within an appropriate sensitivity range, and in intact tissues that retain regular vascular reactivity^8,10,11^. However, no such methods are available for experimental animals, let alone human patients. Implanted devices only record the partial pressure of O_2_ (PO_2_) at a particular spot. Clark-type polarographic electrodes are typically used to measure PO_2_ in blood samples. Although miniaturized for the direct measurement of tissue oxygenation, they are highly invasive. The same is true for fiber-optic probes (or optical electrodes) with O_2_-sensitive luminescent dyes at the tip. Tissue intactness is crucial for oxygenation measurement because contamination by blood cells and plasma results in a spurious increase in the obtained PO_2_. In addition, body intactness is important for systemic regulation studies because surgery substantially affects hemodynamics. By contrast, near-infrared spectroscopy (NIRS) noninvasively monitors O_2_ saturation in blood hemoglobin (Hb). Similarly, the blood O_2_ level-dependent (BOLD) technique, which is based on magnetic resonance imaging, is also available. However, blood Hb-O_2_ saturation does not reflect tissue O_2_ levels. Even if it did, it could only estimate tissue hypoxia by blood desaturation.

One solution to this problem was uncovered through a discovery that we made in an experiment using an all-engineered bioluminescence imaging (BLI) system, AkaBLI^12^. Owing to its highly deliverable and stable luciferin analog (AkaLumine) and its near-infrared emission, which can penetrate most animal tissues and bodies, AkaBLI enables noninvasive visualization of luciferase (Akaluc)-expressing cells deep inside experimental animals. We discovered that the AkaBLI signal from anesthetized mouse striatum was considerably weakened when the animal was accidentally choked during compulsory oral administration, and that the signal shot up when the animal subsequently received O_2_ administration for emergent therapy. Leveraging AkaBLI, we propose a new technology platform, OxyBLI, for the direct measurement of cellular oxygenation in intact experimental animals, mostly rodents. In the present study, which involved hypoxic and/or hyperoxic conditions, the bioluminescence produced by Akaluc’s catalysis of AkaLumine was considered an OxyBLI signal. Mice are genetically manipulatable for their OxyBLI signal formation patterns. On the other hand, rats are more suitable for physiological experiments; their respiro-circulatory system can be well controlled and monitored in parallel with OxyBLI. We found that cellular oxygenation responses in crisis situations were so dynamic in space and time that these could not be detected by any other conventional approaches.

## Results

### Cellular O_2_ levels under hypoxic and hyperoxic environments across mouse organs and cell types

We used the pAAV2 SynTetOff system^13^ to express the Venus-Akaluc fusion protein (Supplementary Fig. 1A) in neurons of the mouse striatum. Forty weeks after viral infection, we anesthetized a spontaneously breathing mouse with isoflurane for BLI. After the intraperitoneal (ip) administration of the hydrochloride salt of AkaLumine (AkaLumine-HCl), a strong and long-lasting bioluminescent signal was observed over the head (Fig. 2A, left). The baseline of the OxyBLI signal was nearly flat approximately two hours after the ip administration. Since Venus-Akaluc is localized inside cells, an OxyBLI signal is hereinafter referred to as the cellular O_2_ level (COL). Significant changes in COL were observed when the fraction of inspired O_2_ (FiO_2_) was changed rapidly between normoxia and hypoxia (Fig. 2A, right; Supplementary Video 1). Hypoxia at 0.1 FiO_2_ reduced COL to half, and severe hypoxia at 0.05 FiO_2_ reduced COL to a negligible level. The COL overshot the baseline value markedly after each hypoxia, with the peak of the overshoot being larger for the FiO_2_ return from 0.05 than that from 0.1.

**Figure 2.**
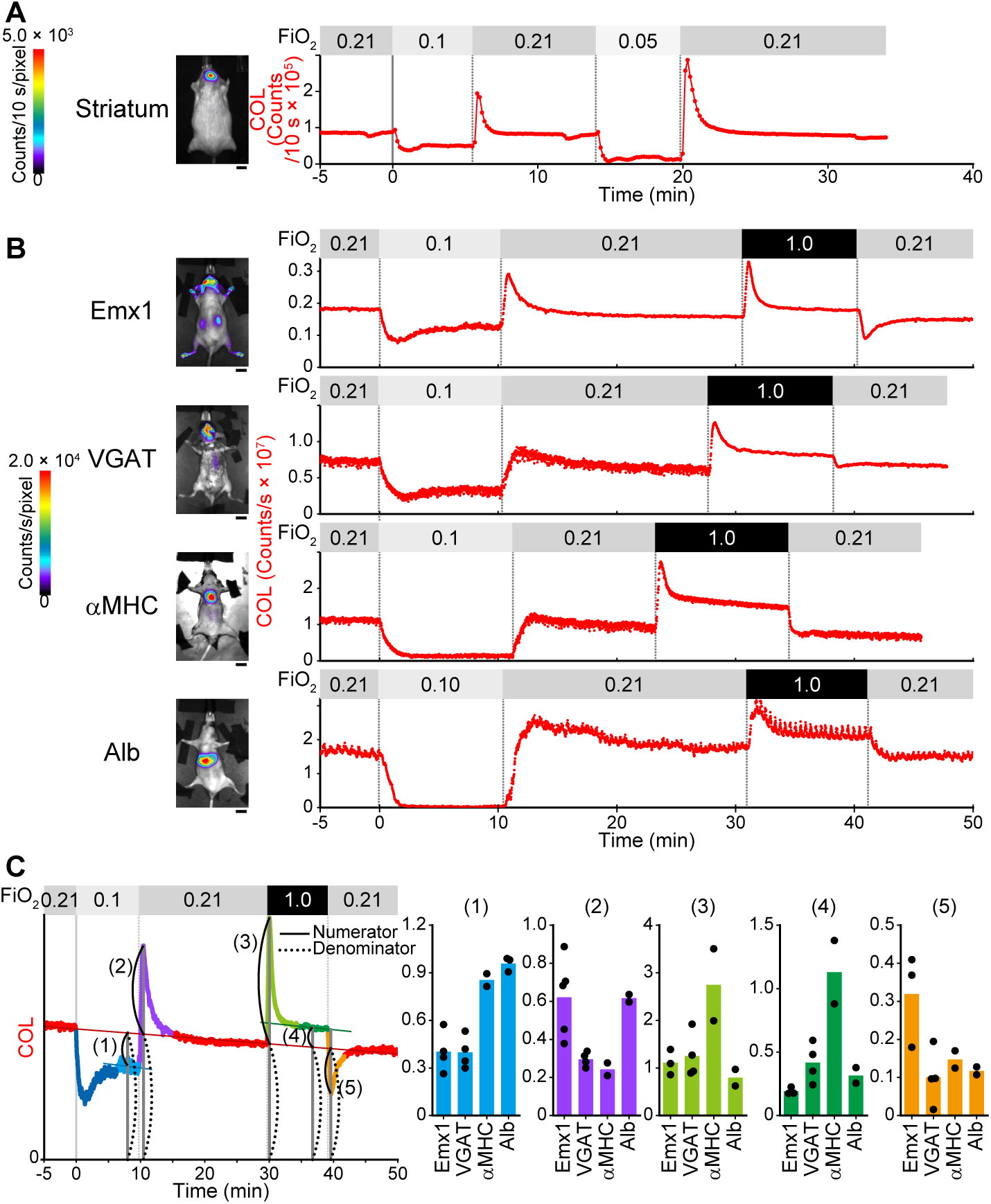
*In vivo* OxyBLI experiments using spontaneously breathing mice under hypoxic and hyperoxic environments. **A,** Time course of striatal COL (cellular O_2_ level) when FiO_2_ was changed between normoxia and hypoxia. The color bar indicates bioluminescence counts/10 s/pixel. **B,** Time courses of COLs in excitatory (Emx1) and inhibitory (VGAT) neurons in the brain, cardiomyocytes in the heart (αMHC), and hepatocytes in the liver (Alb) under hypoxic and hyperoxic conditions. Representatives of *n* = 2–5 repetitions. The color bar indicates bioluminescence counts/s/pixel. **C,** Kinetic analysis of COL changes observed in (**B**). *left,* Five indicators calculated on the basis of the Emx1 COL time course. The numerators (solid curves) were measured with respect to the baseline (straight red line) level: the decrease in the steady state (1); the maximum increase in the overshooting peak (2 and 3); the increase in the steady state (4); and the maximum decrease in the undershooting peak (5). The baseline levels were used as the denominator (dotted curves). *right,* Indicator values are plotted (black dots), and mean values are shown by colored bars. **A,B,** Scale bars: 10 mm. **A–C,** The fraction of inspired O_2_ (FiO_2_) is indicated by shades of gray.

Next, we harnessed the Cre/lox system for cell type-specific control of transgene expression in mice. We recently generated a floxed mouse strain that has a Cre-dependent reporter construct carrying Venus-Akaluc in the ROSA26 locus^14^. By crossing this strain with Emx1-Cre^15^, VGAT-Cre^16^, αMHC-Cre^17^, and Alb-Cre^18^ drivers, we localized Venus-Akaluc to the excitatory neurons of the dorsal forebrain, inhibitory neurons of neural tissues, cardiomyocytes, and hepatocytes, respectively (Supplementary Figs. 1B–E). After the ip administration of AkaLumine-HCl, the brains of Emx1-Venus-Akaluc and VGAT-Venus-Akaluc mice, the heart of αMHC-Venus-Akaluc, and the liver of Alb-Venus-Akaluc mice, produced strong bioluminescent signals (COLs) of similar radiances (Fig. 2B, left). These mice exhibited several characteristic changes in COL when subjected sequentially to a 10-min hypoxia at 0.1 FiO_2_, a recovery period (10–20 min) in room air (0.21 FiO_2_), and a 10-min hyperoxia at 1.0 FiO_2_ (Fig. 2B, right; Supplementary Video 1). First, under the hypoxic condition, the COLs in the brain slightly undershot the target values, which were approximately two-thirds and half of their baseline levels in the excitatory and inhibitory neurons, respectively. On the other hand, the COLs in the heart and liver fell to marginal and imperceptible levels, respectively. These differences suggest compensatory adjustments against acute systemic hypoxia, which include preferential redistribution of blood flow to the brain. Second, when FiO_2_ was returned from 0.1 to 0.21, a salient overshoot was seen in the excitatory neurons, whereas only modest overshoots were noted in the other cell types. Third, upon the onset of hyperoxia, all the cell types produced appreciable overshoots. Fourth, under the hyperoxic condition, the increased steady-state value of COL relative to the baseline value was higher in cardiomyocytes than in the other cell types. Fifth, when FiO_2_ was returned from 1.0 to 0.21, an apparent undershoot was observed in the excitatory neurons but not in the other cell types. These five measurements were repeated using other mice, and similar results were obtained (Fig. 2C). The heterogeneity in COL dynamics at the organ level may be mainly explained by the systemic regulation of blood flow. On the other hand, the difference in COL dynamics detected within an organ may be explained by the local difference in the O_2_ demand–supply balance. It is possible that the balance differs between excitatory and inhibitory neurons in the brain. The O_2_ demand is likely higher in inhibitory neurons, whose discharge plays an important role in high-frequency network activities, than in excitatory neurons^19^. Thus, an excessive consumption of O_2_ in inhibitory neurons may have resulted in the absence of a salient overshoot of COL at the end of the hypoxia at 0.1 FiO_2_. Additionally, excitatory neurons may be more sensitive to hyperoxia than inhibitory neurons. This may account for the fact that the COL overshoots in excitatory neurons had shorter lifetimes than those in inhibitory neurons and that the transient hyperoxia (1.0 FiO_2_) was followed by a substantial undershoot in excitatory neurons but not inhibitory neurons.

### *Ex vivo* O_2_ titration using detached mouse auricles

The *in vivo* performance of OxyBLI motivated us to examine how Akaluc depends on O_2_ in comparison with the conventional firefly luciferase (Fluc). Because of its hydrophobic nature, however, O_2_ does not diffuse well through the static layer of aqueous medium overlaying cultured cell monolayers^11^. Thus, conventional culture experiments cannot replicate the *in vivo* O_2_ delivery regulation. Conventional *in vitro* experiments that employ aqueous solutions face similar limitations. Accordingly, it is challenging to measure the O_2_ dependency of bioluminescence, which primarily functions in hydrophilic environments. Aiming at the airborne application of O_2_, we took an interest in the mouse auricle. The very thin part of the auricle has a large surface area. We found that the auricle of an Emx1-Venus-Akaluc mouse emitted strong bioluminescence signals (Fig. 3A). Histological studies of auricular transverse sections revealed the localized Venus-Akaluc fusion protein expression in hair follicles, sebaceous glands, and basal keratinocytes (Supplementary Fig. 2A, left). Assuming that Akaluc would respond to O_2_ blown from the outside, we expected a detached auricle to serve as an *ex vivo* device for titrating Akaluc with different O_2_ concentrations. Dissection results in a loss of blood supply and neural regulation in the auricle, which would allow for straightforward interpretation of the device’s signals.

**Figure 3.**
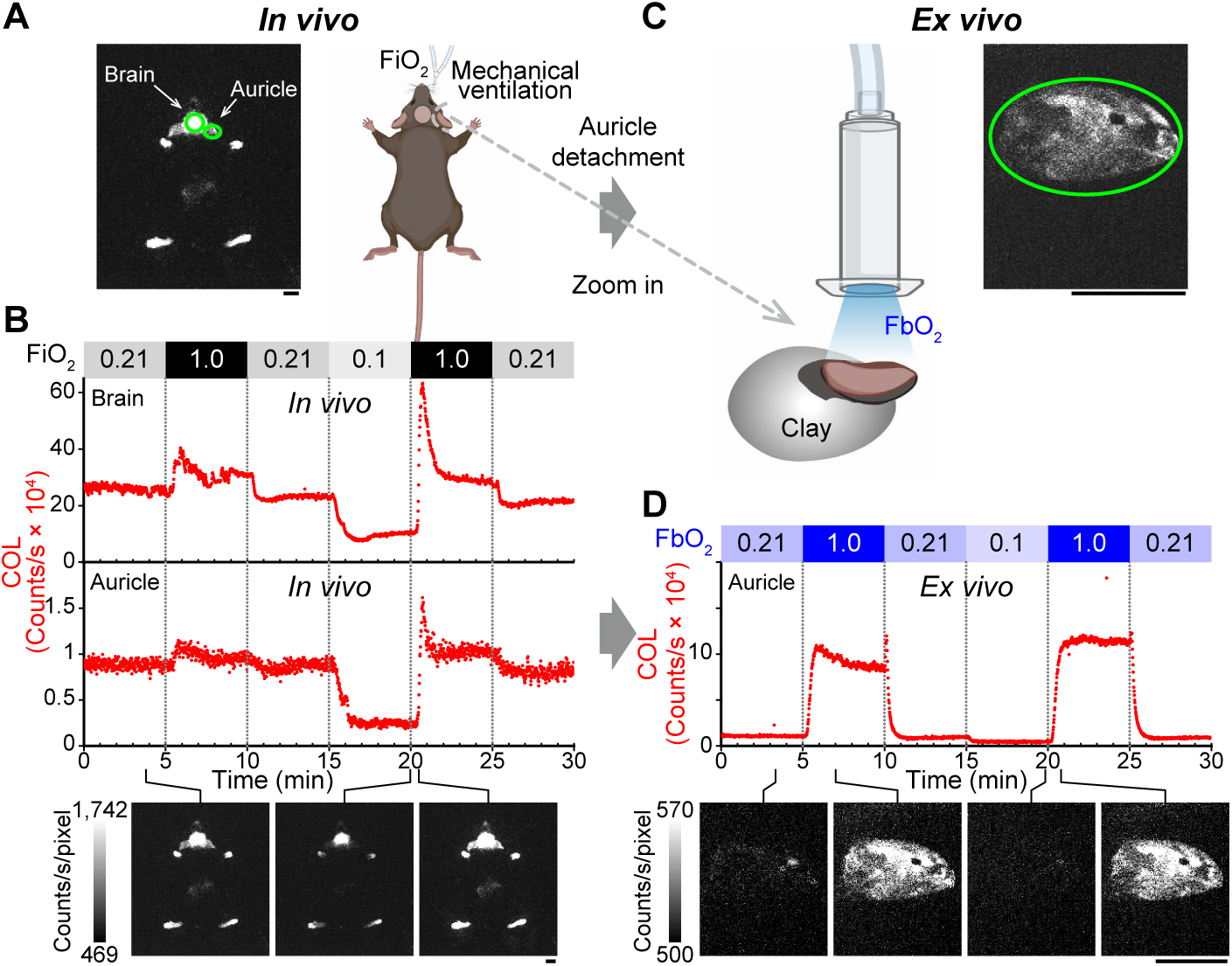
Successive *in vivo*/*ex vivo* OxyBLI experiments using mouse auricles. An Emx1-Venus-Akaluc mouse was used. Right-pointing gray arrows indicate the transfer of BLI from *in vivo* to *ex vivo*. Scale bars: 10 mm. **A,B,** *In vivo* OxyBLI experiment. **A,** *left,* A bioluminescence (BL) image of the intubated mouse after ip administration of AkaLumine-HCl. *right,* A schematic of an *in vivo* imaging experiment, in which the fraction of inspired O_2_ (FiO_2_) was changed. **B,** Time courses of COLs in the brain and the right auricle during the regular FiO_2_ protocol, which included a 5-min hyperoxia at 1.0 FiO_2_ and a 5-min hypoxia at 0.1 FiO_2_ followed by a 5-min hyperoxia at 1.0 FiO_2_ with 0.21 FiO_2_ as baseline. **C,D,** *Ex vivo* OxyBLI experiment. **C,** After detachment, the right auricle was placed on clay and subjected to an *ex vivo* imaging experiment that used a close-up lens. *right,* A BL image zooming in on the detached auricle. *left,* A schematic of an *ex vivo* imaging experiment, in which the fraction of blown O_2_ (FbO_2_) was changed. **D,** Time course of COL in the detached auricle during the regular FbO_2_ protocol, which included a 5-min hyperoxia at 1.0 FbO_2_ and a 5-min hypoxia at 0.1 FbO_2_ followed by a 5-min hyperoxia at 1.0 FbO_2_ with 0.21 FbO_2_ as baseline. **B,D,** Shown below the graphs are representative BL images at indicated times. The gray scale indicates the lowest and highest intensities of the image. FiO_2_ and FbO_2_ are indicated by shades of gray and blue, respectively.

To investigate the dynamic interplay between hypoxia and hyperoxia in this study, we designed a regular FiO_2_ protocol that included a 5–10-min hyperoxia at 1.0 FiO_2_ (1^st^ phase) and a 5–10-min hypoxia at 0.1 FiO_2_ followed by a 5–10-min hyperoxia at 1.0 FiO_2_ (2^nd^ phase) with 0.21 FiO_2_ as the baseline. We first performed an *in vivo* OxyBLI experiment with the regular FiO_2_ protocol that included 5-min hypoxia and hyperoxia using an Emx1-Venus-Akaluc mouse to monitor signals from the brain and auricle (Figs. 3A and 3B; Supplementary Video 2, left). The COL response to hyperoxia (1.0 FiO_2_) was characterized again by an overshoot followed by a steady-state phase, suggesting the occurrence of a vasoconstrictor response in both organs. Next, we separated the right auricle at the base and placed it on clay to expose it to blowing air at a velocity of 4 L/min (Fig. 3C). For this *ex vivo* OxyBLI, we used a close-up lens to zoom in on this object and changed the fraction of blown O_2_ (FbO_2_) in the same way as the regular FiO_2_ protocol (Fig. 3D; Supplementary Video 2, right). At 1.0 FbO_2_, the COL increased markedly and remained at a high level. These results strongly suggest that Akaluc in tissues is supplied with excess amounts of ATP and AkaLumine to produce emission exclusively in an O_2_-dependent manner. We performed similar successive *in vivo*/*ex vivo* experiments while keeping the distance between the auricle and the lens constant (Supplementary Fig. 2A, right). This imaging setup allowed for a direct comparison of COL intensities between *in vivo* and *ex vivo* situations; we found that the *ex vivo* baseline was approximately one order of magnitude lower than the *in vivo* baseline. We also employed CAG-Venus-Akaluc mice (Supplementary Fig. 2B) and observed the same FbO_2_-dependent changes in auricular COL as for Emx1-Venus-Akaluc mice. To examine the O_2_ dependency of Fluc, furthermore, we employed CAG-*ff*Luc-cp156 transgenic mice^20^, which produce *ff*Luc-cp156, a chimeric protein containing Fluc and a circularly permuted Venus. We prepared auricle samples from CAG-Venus-Akaluc and CAG-*ff*Luc-cp156 mice after administering the respective luciferins to perform side-by-side comparison experiments. Whereas a similar O_2_ dependency was observed between the two luciferases, the dynamic range of Akaluc was one order of magnitude larger than that of Fluc (Supplementary Fig. 3).

### Cellular O_2_ levels in rat brain with systemic parameters under hypoxic and hyperoxic environments

We next monitored macrovascular hemodynamics and basic vital signs with striatal OxyBLI signals simultaneously to clarify the dynamic effects of FiO_2_ changes on brain O_2_ levels in a systemic context. We established a rat model for a multi-sided approach to whole-body monitoring. The configuration of the experimental setup with a rat is illustrated in Figure 4A. Skin-retracted rat heads were imaged (COL). Regional cerebral blood flow (CBF) was measured in the left striatum with a laser Doppler flowmeter. The left and right femoral arteries were equipped with arterial catheters for measuring PO_2_ in arterial blood (PaO_2_) and blood pressure (BP), respectively. Heart rate (HR) and respiratory rate (RR) were monitored through an electrocardiogram (ECG). SpO_2_ was monitored by attaching a pulse oximeter to the right forelimb. OxyBLI experiments started a few hours after AkaLumine-HCl ip administration. In one experiment, we used a single rat with striatal bioluminescence over the head to perform the following three protocols for the FiO_2_ challenge.

**Figure 4.**
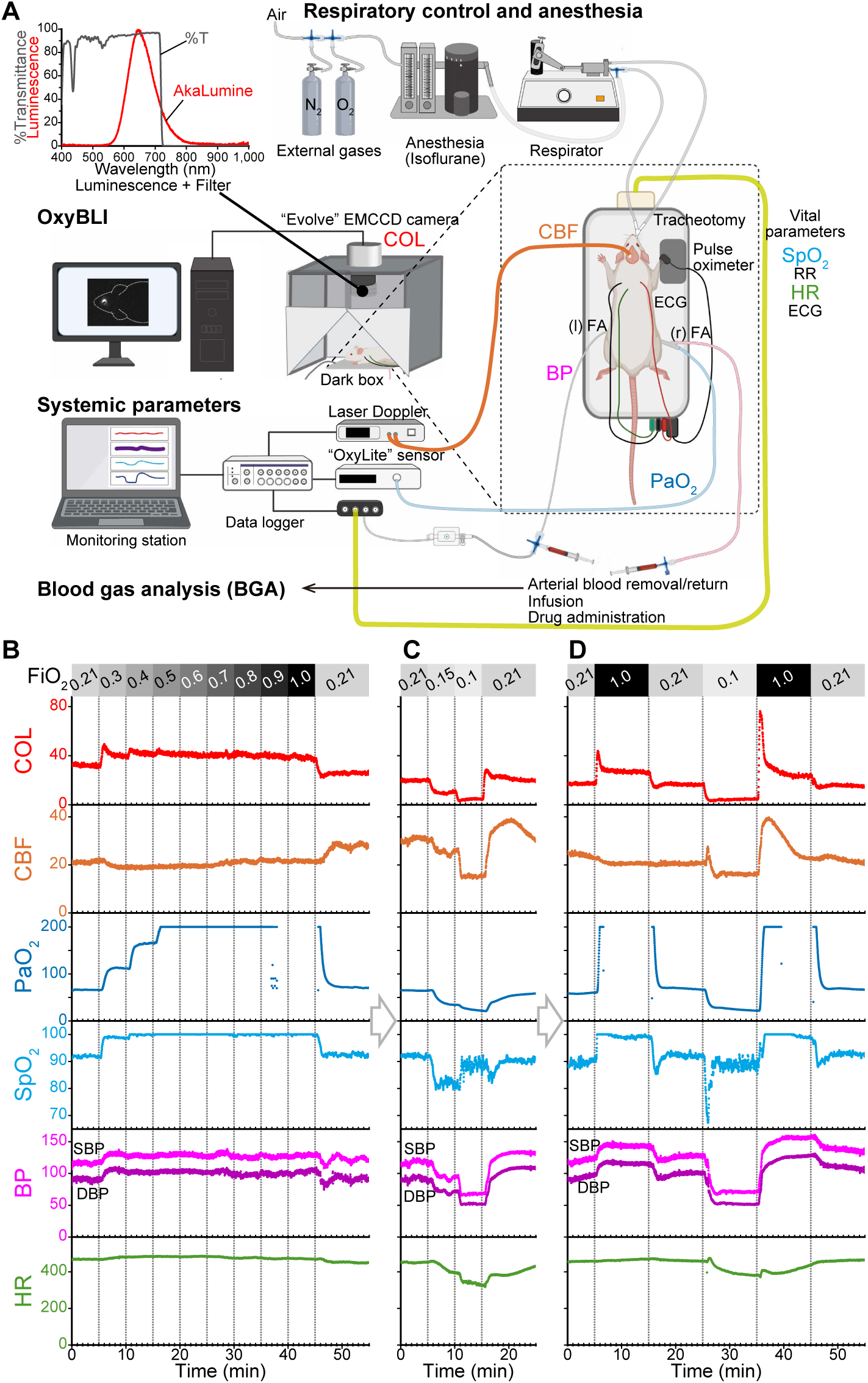
Understanding the dynamics of rat brain tissue oxygenation in a systemic context. A rat expressing Venus-Akaluc in the striatum was intubated for artificial ventilation and isoflurane anesthesia. Measured physiological indicators are as follows. COL (red): cellular O_2_ level, CBF (orange): cerebral blood flow, PaO_2_ (blue): partial pressure of arterial O_2_, SpO_2_ (light blue): saturation of percutaneous O_2_, BP (magenta): blood pressure, and HR (green): heart rate. **A,** Configuration of the *in vivo* experimental system. The 780-nm light from the laser Doppler flow cytometer for CBF measurement was eliminated by three short-pass filters that can pass most of the AkaLumine emission. A data logger was used to record and monitor the systemic parameters continuously. Abbreviations: RR: respiration rate; ECG: electrocardiogram; FA: femoral artery. **B–D,** Time courses of striatal COL and systemic parameters during three types of FiO_2_ challenges: stepwise increases from 0.21 to 1.0 (**B**), graded decreases from 0.21 to 0.1 (**C**), and the regular protocol that included 0.1 FiO_2_ and 1.0 FiO_2_ (**D**). The fraction of inspired O_2_ (FiO_2_) is indicated by shades of gray. These experiments, shown in (**B–D**), were conducted sequentially (right-pointing open arrows) on a single rat. Similar results were obtained from two other independent experiments that observed rat striatal COL. Under the respiratory support illustrated (blood gas analysis). Similar PaO_2_ values were observed in the traces obtained by the OxyLite sensor, which was inserted into the left femoral artery. Notably, however, this sensor has a maximum measurement range of 200 mmHg. Abbreviations: SBP: systolic BP. DBP: diastolic BP.

First, we gradually increased FiO_2_ from 0.21 to 1.0 in a stepwise fashion (Fig. 4B). The initial increase from 0.21 to 0.3 resulted in substantial increases in PaO_2_ and SpO_2_. This stimulus probably resulted in vasoconstriction, which led to an increase in BP and a decrease in CBF. An elevation of striatal COL signal suggested that the increase in blood O_2_ level surpassed the decrease in CBF. The signal elevation was composed of a modest overshoot and a subsequent new steady-state level. The second FiO_2_ increase from 0.3 to 0.4 resulted in a smaller overshoot and a slightly higher steady-state level, which proved to be saturable because it was not affected by further FiO_2_ increases of up to 1.0. After the whole challenge was over, all signals returned to pre-stimulus levels observed at 0.21 FiO_2_.

Second, we decreased FiO_2_ in two steps. The rat was exposed to two consecutive 5-min hypoxic states at 0.15 and 0.1 FiO_2_ (Fig. 4C). Possible hypoxia-induced parasympathetic activation decreased HR and, consequently, CBF in both steps. Such a decrease in blood flow, in conjunction with the expected decrease in PaO_2_, reduced the COL signal linearly. However, when FiO_2_ was returned from 0.1 to 0.21, the COL signal returned quickly with a modest overshoot, whereas CBF recovered slowly.

Third, we performed the regular FiO_2_ protocol (see Fig. 3B) that included 10-min hypoxia and hyperoxia, and observed typical changes in COL signals (Fig. 4D; Supplementary Video 3). Notably, clear overshoots appeared at the onset of 1.0 FiO_2_. When FiO_2_ increased from 0.21 (1^st^ phase), CBF fell slightly and the COL overshoot was mild. However, when FiO_2_ increased from 0.1 (2^nd^ phase), CBF increased transiently but to a great extent, and the overshoot increased by approximately threefold compared to the 1^st^ phase. In another experiment, the regular protocol was followed by an additional FiO_2_ challenge, which involved changing from a 10-min hypoxia to a 10-min hyperoxia through a 5-min normoxia (Fig. 5). This two-step transition suppressed the overshooting magnitude in both steps. It was therefore concluded that the magnitude of the hyperoxia-induced COL overshoot depended on the preceding COL. After overshooting, however, the COL signals under hyperoxic conditions (1.0 FiO_2_) gradually reached steady-state values that were nearly identical between the 1^st^ and 2^nd^ phases (Figs. 4D and 6A). We repeated this experiment, testing the regular FiO_2_ protocol on different rat samples (Fig. 6B). The increase in overshooting magnitude in the 2^nd^ phase compared to the 1^st^ phase was statistically significant. In contrast, the eventual increments compared to the baseline at 0.21 FiO_2_ were nearly identical: 41.4% ± 6.6% (*n* = 6) and 44.1% ± 8.6% (*n* = 6) for the 1^st^ and 2^nd^ phases, respectively. These results suggest that the steady-state level during hyperoxic breathing remains consistent regardless of the preceding conditions.

**Figure 5.**
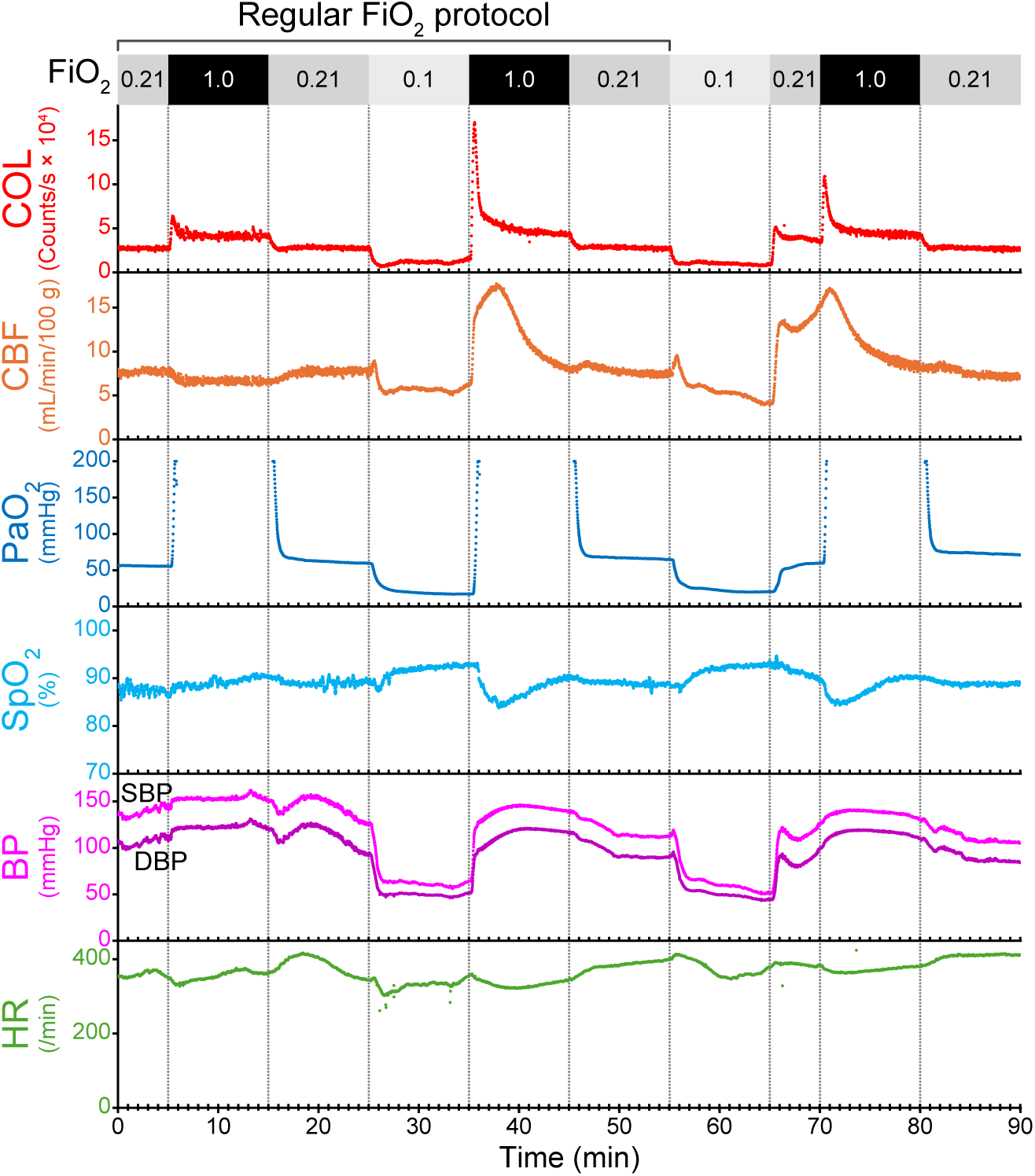
Magnitude of the hyperoxia-induced COL overshoot. Time courses of striatal COL and systemic parameters during the regular FiO_2_ protocol followed by an additional FiO_2_ challenge with changes from a 10-min hypoxia (0.1 FiO_2_) to a 10-min hyperoxia (1.0 FiO_2_) through a 5-min normoxia (0.21 FiO_2_). A rat expressing Venus-Akaluc in the striatum was intubated for artificial ventilation and isoflurane anesthesia. The experimental setup was identical to that shown in Figure 4A.

**Figure 6.**
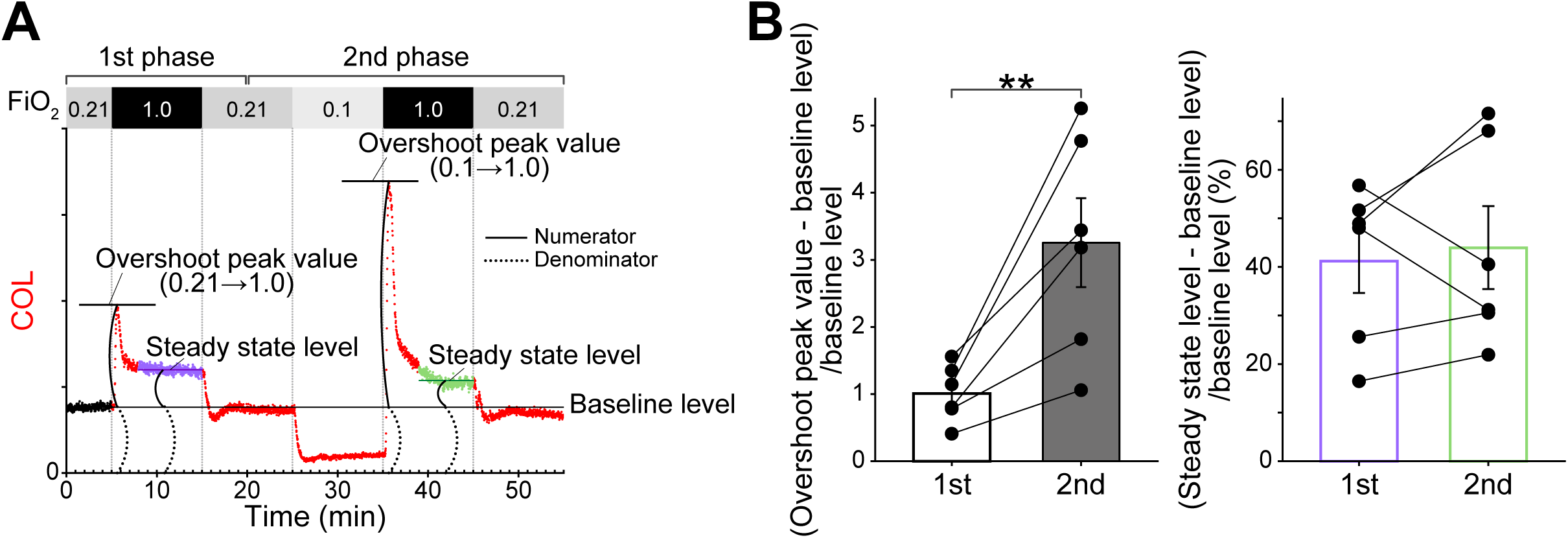
Hyperoxia-induced changes in COL in the transition and steady states. **A,** The regular FiO_2_ protocol uses 0.21 FiO_2_ as the baseline and consists of a first phase that includes 10 minutes of hyperoxia at 1.0 FiO_2_ and a second phase that includes 10 minutes of hypoxia at 0.1 FiO_2_ followed by 10 minutes of hyperoxia at 1.0 FiO_2_. Two indices: (overshoot peak value – baseline level)/baseline level and (steady state level – baseline level)/baseline level, were calculated. These reflect to hyperoxia-induced changes in COL in the transition and steady states, respectively. Their calculation is illustrated based on the time course shown in Figure 4D. In principle, the numerators (solid curves) were measured with respect to the baseline level, which was used as the denominator (dotted curves). **B,** Statistics of the two aforementioned indices are shown. The regular FiO_2_ protocol was performed in six independent experiments, including those that produced Figures 4D, 5 and 8C. The difference between the first and second phases was significant for the overshooting magnitude (left), but not for the steady-state level (right). The mean values ± s.e.m. (*n* = 6 rats) are shown by bars. Statistical significance (** *P* < 0.01) was examined by Welch’s unpaired *t* test.

### Respiratory control

Hyperventilation lowers arterial carbon dioxide pressure (PaCO_2_). Hypocapnia, defined as a PaCO_2_ below 35 mmHg, results in vasoconstriction of the cerebral arterioles and reduces cerebral blood flow and volume, thereby decreasing intracranial pressure. Thus, controlled mechanical hyperventilation is used in neurointensive care to treat intracranial hypertension in traumatic brain injury patients^21,22^. However, the benefits of intraoperative hyperventilation remain controversial owing to a lack of information on brain oxygenation. To examine how brain hypoxia develops during hyperventilation, we temporarily increased RR in an intubated rat expressing Venus-Akaluc in the right striatum (Fig. 7A, left). As the conventional mode of mechanical ventilation during surgery involves slightly increasing FiO_2_ to 0.3–0.5 (refs. 23 and 24), we set FiO_2_ at 0.5. When RR increased from 50 to 100 and then to 125, CBF decreased in a gradient manner, and striatal COL decreased in the same manner. We performed blood gas analysis (BGA) intermittently and found that PaO_2_ increased only slightly, whereas PaCO_2_ was within the expected range for hypocapnia during the hyperventilation.

**Figure 7.**
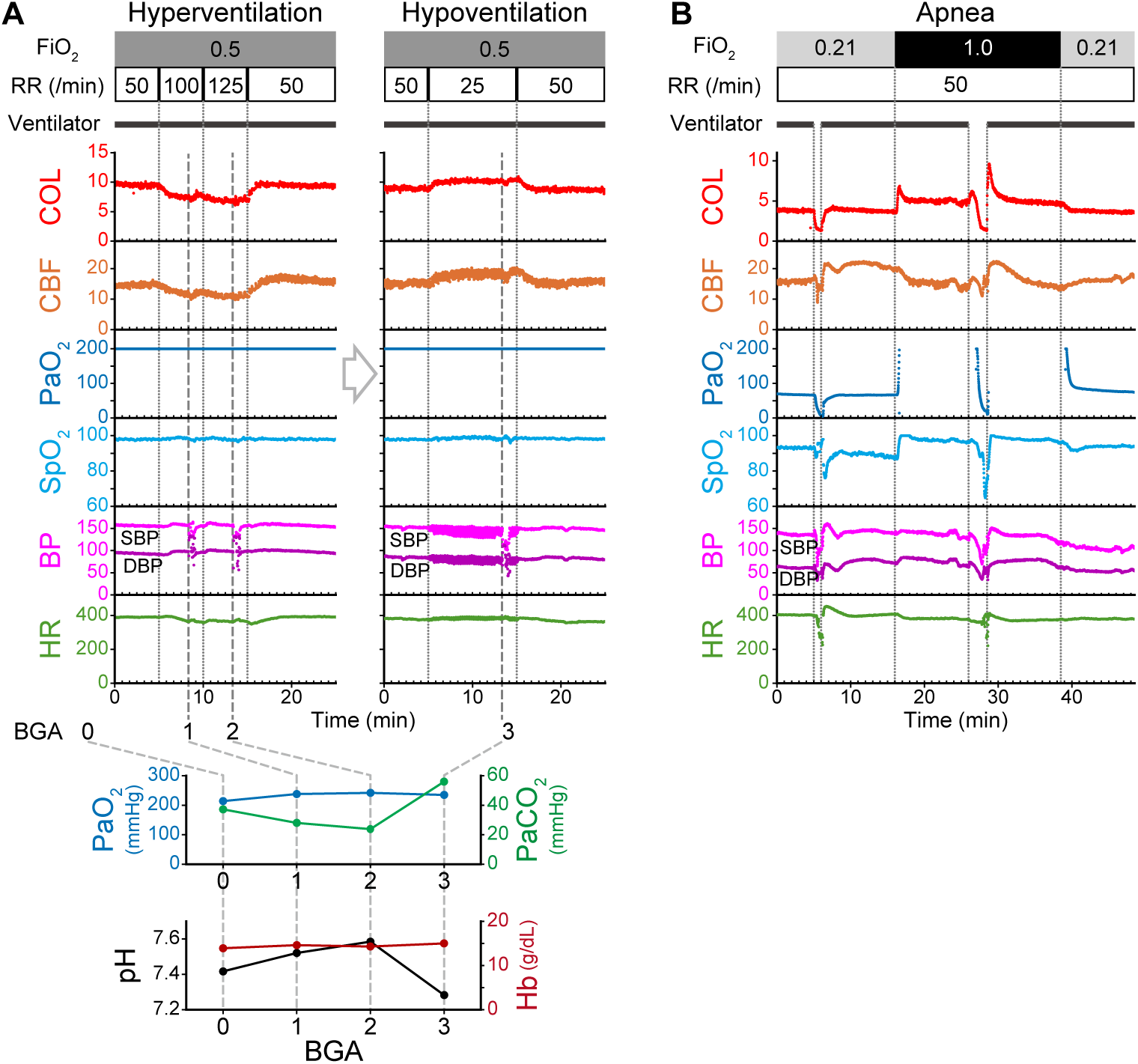
Cellular O_2_ levels in the rat striatum during modified ventilatory patterns. Rats expressing Venus-Akaluc in the striatum were intubated for artificial ventilation and isoflurane anesthesia. Measured physiological indicators are as follows. COL (red): cellular O_2_ level, CBF (orange): cerebral blood flow, PaO_2_ (blue): partial pressure of arterial O_2_, SpO_2_ (light blue): saturation of percutaneous O_2_, BP (magenta): blood pressure, and HR (green): heart rate. Abbreviations. RR: respiratory rate. SBP: systolic BP. DBP: diastolic BP. The experimental setup was identical to that shown in Figure 4A. The fraction of inspired O_2_ (FiO_2_) is indicated by shades of gray. **A,** *left,* Hyperventilation. *right,* Hypoventilation. These two experiments were conducted sequentially (indicated by a right-pointing open arrow) on a single rat. *bottom,* Blood gas analysis (BGA) data obtained immediately before the experiment (“0”) and at times indicated by “1”, “2”, and “3”. Representative of *n* = 3 independent experiments using different rats. **B,** Apnea. Representative of *n* = 2 independent experiments using different rats.

Conversely, both CBF and striatal COL increased in a controlled mechanical hypoventilation experiment, wherein RR was reduced from 50 to 25 (Fig. 7A, right). BGA confirmed that the hypoventilation caused hypercapnia (PaCO_2_ = 56.0 mmHg) and respiratory acidosis (pH = 7.283), but only a slight change in PaO_2_. ‘Permissive hypercapnia’ is a ventilatory strategy for treating acute respiratory failure^25^, and is expected to protect the lungs through ventilation with low inspiratory volume and pressure while maintaining gas exchange. An additional benefit of permissive hypercapnia is improved tissue oxygenation. Monitoring COL in the brain will be useful in assessing the validity of permissive hypercapnia in various life-threatening situations in experimental animals.

Lastly, we subjected the same rat to apnea at 0.21 FiO_2_ and 1.0 FiO_2_ (Fig. 7B). Apnea was induced by stopping the ventilator; as soon as bradycardia was noted, ventilation resumed. As PaO_2_ gradually approached zero during apnea, striatal COL also decreased substantially under both FiO_2_ conditions. At 1.0 FiO_2_, however, the onset of apnea was accompanied by small, transient increases in COL and CBF. These transient increases were reproducibly observed at 1.0 FiO_2_ but not at 0.21 FiO_2_, suggesting that the microvascular system is hypersensitive to decreases in PaO_2_ when exposed to hyperoxia.

### Bleeding and autotransfusion

Using the arterial line in the rat model system (Fig. 4A), we performed a blood salvage experiment while observing striatal OxyBLI signals. The experiment was composed of a bleeding phase, during which 2.5 mL of blood was collected three times at 5-min intervals and a recovery phase, during which the collected blood (7.5 mL) was transfused into the animal 5 min after the last collection (Fig. 8A, left). These manipulations significantly altered BP in the femoral artery. To understand how COL in the striatum buffered flow (CBF) and pressure (mBP: mean BP), we constructed correlation plots (Fig. 8A, right) between mBP and CBF, CBF and COL, and mBP and COL. During the bleeding phase, mBP decreased considerably, as expected. First, a conventional pressure–flow relationship was apparent in the mBP–CBF plot, suggesting the existence of classic ‘cerebral autoregulation’^26^. CBF decreased only slightly when mBP was reduced to 55 mmHg from a baseline of 90 mmHg and became pressure-passive at mBP < 55 mmHg. The former part may reflect a plateau, and the mBP value at the inflection point (55 mmHg) may represent the lower limit. Second, in the CBF–COL plot, COL was dependent on CBF below 6 mL/min/100 g; the data points are densely scattered over a CBF range of 6–9 mL/min/100 g. Lastly, the mBP–COL plot may be the product of the mBP–CBF and CBF–COL plots. Like CBF, COL was pressure-passive at mBP < 55 mmHg but not so within the mBP range above the lower limit (55–90 mmHg).

**Figure 8.**
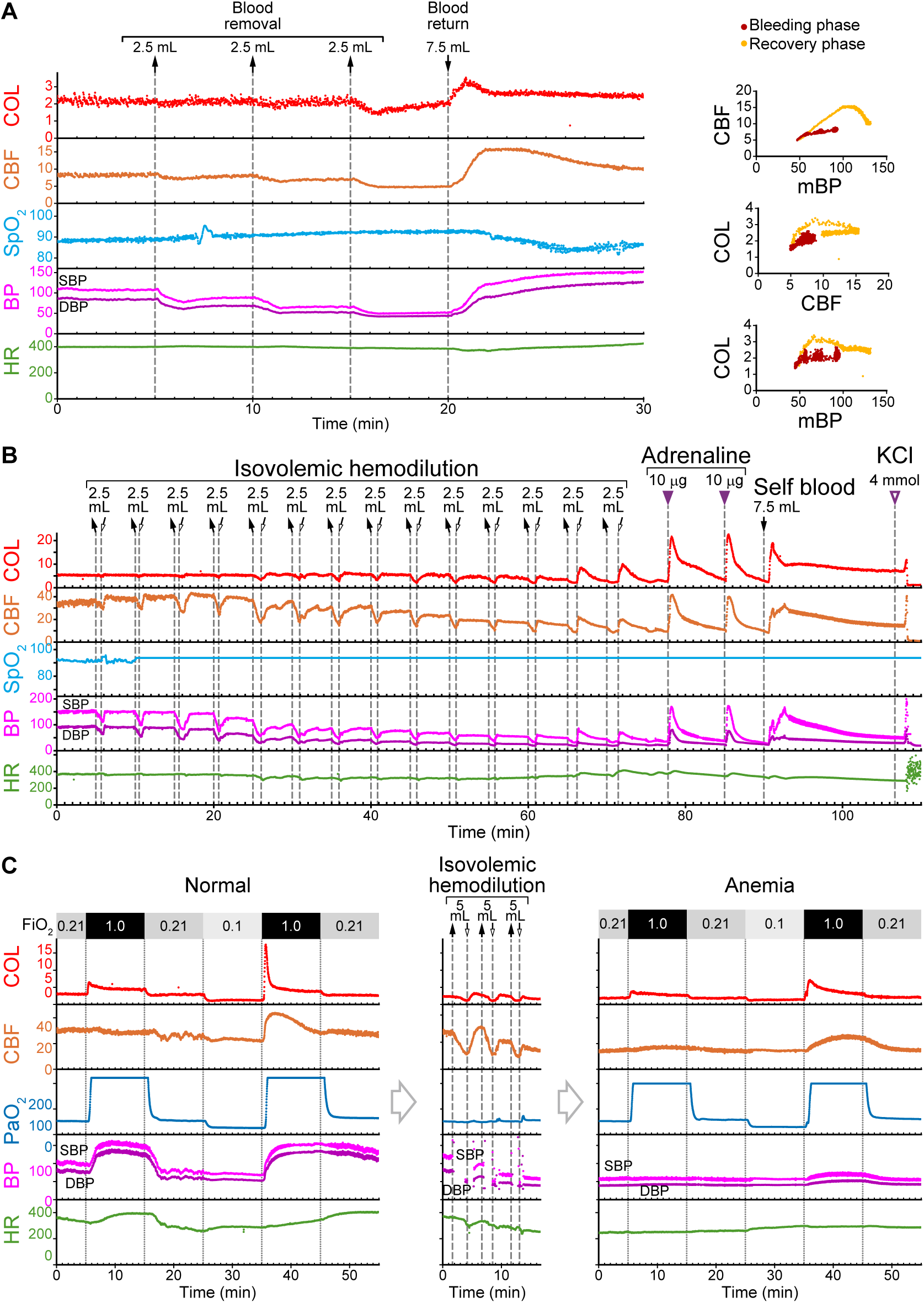
Cellular O_2_ levels in the rat striatum in the anemic state. Rats expressing Venus-Akaluc in the striatum were intubated for artificial ventilation and isoflurane anesthesia. Measured physiological indicators are as follows. COL (red): cellular O_2_ level, CBF (orange): cerebral blood flow, PaO_2_ (blue): partial pressure of arterial O_2_, SpO_2_ (light blue): saturation of percutaneous O_2_, BP (magenta): blood pressure, and HR (green): heart rate. Abbreviations. SBP: systolic BP. DBP: diastolic BP. The experimental setup was identical to that shown in Figure 4A. **A,** *left,* Time courses during bleeding and transfusion. *right,* Correlation plots between mBP and CBF (top), CBF and COL (middle), and mBP and COL (bottom). Data points in the bleeding phase (<t = 1,000 sec) are colored red. Data points in the recovery phase (>t = 1,000 sec) are colored yellow. **B,**Time courses while isovolemic hemodilution was followed by resuscitation. Shown is a representative of *n* = 3 independent experiments that involved isovolemic hemodilution. The pulse oximeter failed to measure SpO_2_, likely because of a possible vascular collapse during the hemodilution. **C,** Results of the regular FiO_2_ protocol before and after the isovolemic hemodilution that yielded a final Hb level of 5.1 g/dL. The fraction of inspired O_2_ (FiO_2_) is indicated by shades of gray.

However, COL at mBP > 55 mmHg exhibited substantial fluctuation, suggesting the presence of additional mechanisms that regulate the link between blood flow and O_2_ delivery. Overall, there appears to be a mechanism that maintains COL stability despite marked changes in arterial pressure. During the recovery phase, in contrast, all the plots showed a convex upward trend, and hysteresis was evident in each loop. This hysteresis is considered a lag between mBP and CBF or COL. Although the transfusion was completed in 30 sec, if the process were slowed down, the hysteresis would be less evident.

### Isovolemic hemodilution

Isovolemic hemodilution, a technique in which blood is removed and replaced with isotonic fluids to maintain euvolemia, is used to avert the adverse effects of allogeneic blood transfusions^27,28^. An intubated rat expressing Venus-Akaluc in the right striatum was subjected to repetitive isovolemic hemodilutions through a catheter at the right femoral artery (Fig. 8B). Substitution of 2.5 mL of isotonic saline for the same blood volume occurred at 5-minute intervals. After 14 substitutions, Hb level diminished to a reading below 5.0 g/dL, the lower detection limit of our BGA analyzer. Each infusion improved CBF and striatal COL, thereby corroborating the clinically beneficial effects of this intervention. Notably, COL was well maintained at the baseline level throughout the experimental period. Conversely, the BP and CBF baselines declined substantially. The last two infusions resulted in overshoots in COL, CBF, and BP, suggesting the emergence of hypersensitivity in the macrovascular system in the severely anemic state. In the resuscitation phase, COL, CBF, and BP increased rapidly and decreased gradually over a similar time course upon the administration of adrenaline. Finally, an autologous blood transfusion (7.5 mL) induced a prominent peak and a sustained baseline in COL, suggesting that the O_2_ supply from red blood cells (RBCs) contributes to brain oxygenation in such a critical situation.

Next, we performed the regular FiO_2_ protocol before and after an isovolemic hemodilution (Fig. 8C). BP was almost flat at around 50 mmHg during the anemic state. This unresponsive systemic hypotension indicates altered O_2_-dependent vascular hemodynamics, including CBF regulation. The decrease in magnitude of the 2^nd^ phase overshoot in the anemic state suggested that the O_2_ supply from RBCs contributes to this transient increase in brain oxygenation.

### Monitoring cellular O_2_ levels across multiple organs in crisis situations

Thanks to the homogeneous biodistribution of the AkaLumine substrate, the floxed Venus-Akaluc mouse line is useful for the whole-body characterization of the recombination pattern of any Cre driver system. We noticed that Emx1-Venus-Akaluc mice exhibited substantial bioluminescent signals in the distal portions of the upper and lower extremities and the kidneys^29^ in addition to the strong signals in the brain (Fig. 2B, Emx1). This characteristic pattern was reproducibly identified in all ten Emx1-Venus-Akaluc mice imaged using bioluminescence, and it was corroborated by histological examination of Venus fluorescence (Supplementary Fig. 1B). This multi-site expression of Venus-Akaluc allows for simultaneous monitoring of COLs in the brain, limb muscles, and kidneys. We confirmed that all of the signals were long-lasting. Then, we performed two OxyBLI experiments. First, we focused on the distal localization of Venus-Akaluc in the extremities. This “gloves and socks” distribution led us to use this mouse line as a perfect system for assessing limb tissue reperfusion after arterial occlusion (Fig. 9A). Reactive hyperemia is characterized by quick and excessive reperfusion after a period of ischemia^30^. However, it has been debated whether hyperemia results in tissue hyperoxia. One previous study demonstrated reactive hyperoxia by placing a phosphorescence-based O_2_ sensor on rat hindlimbs^31^. However, it is questionable how reliably the O_2_ quenching method can measure elevated tissue oxygenation levels. We subjected a spontaneously breathing, anesthetized mouse to OxyBLI with a tourniquet on the proximal portion of the left hindlimb. Compression was applied through the tourniquet for 5 min while COLs were monitored in the Emx1-related regions. The COL in the distal portion decreased to nearly zero, indicating efficient occlusion of the hindlimb arteries. Approximately 1 min after the onset of occlusion, RR slightly increased, likely due to a response from the autonomic nervous system. Accordingly, there were slight increases in the COLs in other parts. Significantly, we observed post-occlusion hyperoxia in the left hindlimb muscle cells, and the time course was similar to that of reported reactive hyperemia^31^.

**Figure 9.**
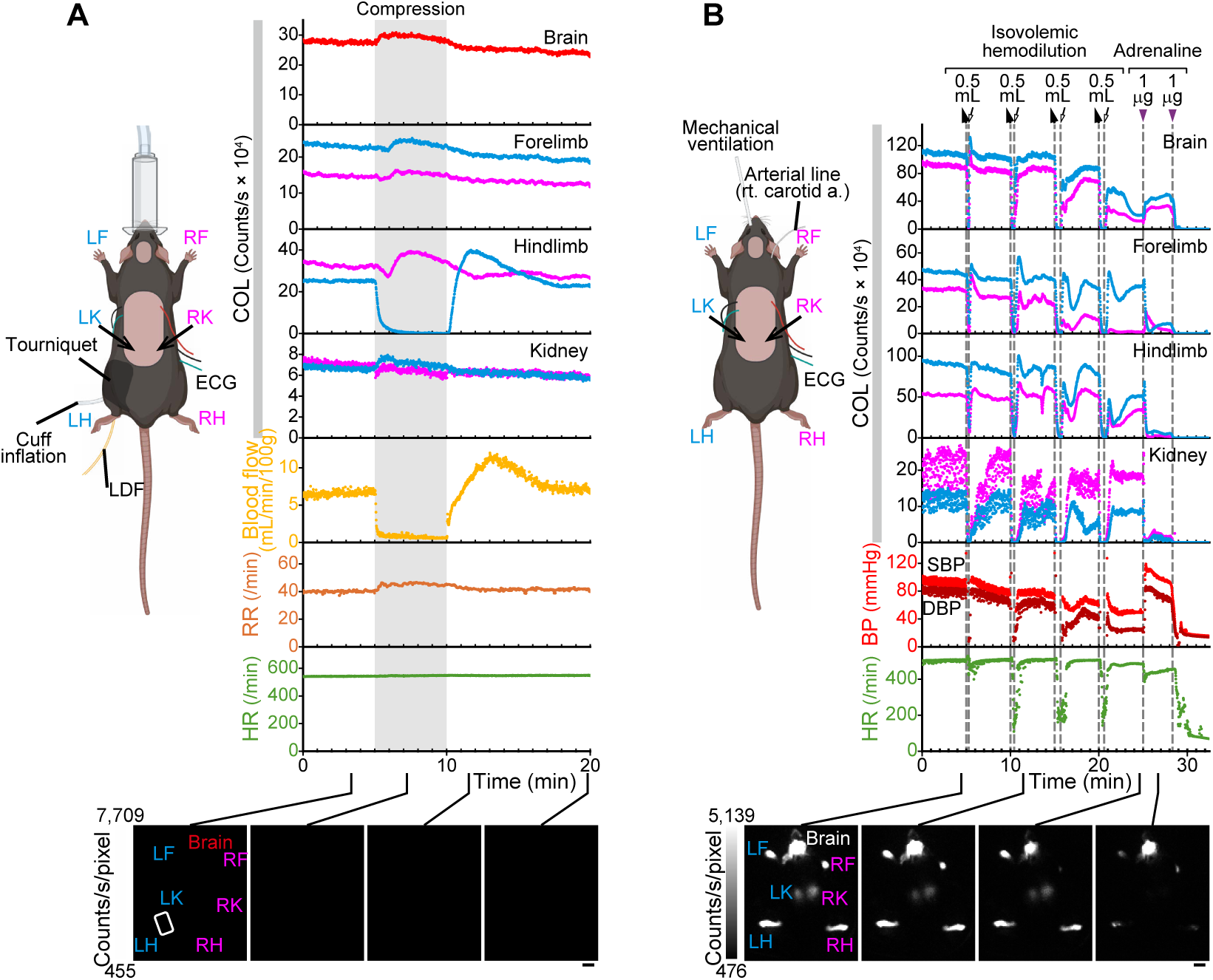
*In vivo* multisite OxyBLI experiments using Emx1-Venus-Akaluc mice. **A,** Post-ischemic reactive hyperoxia, due to a transient increase in blood flow termed reactive hyperemia. A spontaneously breathing mouse was anesthetized with isoflurane. A tourniquet cuff attached to the left thigh was used to stop arterial blood flow into the distal portion (Compression) for 5 min. A laser Doppler flowmeter (LDF) was attached to the heel to measure blood perfusion in the extremity. Respiratory rate (RR) was estimated from the ECG signal. Representative of *n* = 3 experiments that involved compression. **B,** Isovolemic hemodilution and resuscitation. An artificially breathing mouse was anesthetized with isoflurane. A catheter in the right carotid artery was used for hemodilution, BP measurement, and drug administration. Representative of *n* = 2 experiments. **A,B,** Shown below the graphs are representative BL images at indicated times. In each image, the gray scale indicates the lowest and highest intensities of the image. Scale bars: 10 mm. Abbreviations: LF: left forelimb; RF: right forelimb; LK: left kidney; RK: right kidney; LH: left hindlimb; RH: right hindlimb.

For the second experiment, we used another anesthetized mouse and induced acute anemia, followed by cardiopulmonary resuscitation (Fig. 9B). This experiment involved severe treatments; the mouse was intubated for mechanical ventilation. Anemia was induced through four repetitive isovolemic hemodilutions via the right carotid artery. During each hemodilution, 0.5 mL of blood was replaced with 0.5 mL of isotonic saline. The brain COL remained stable and resilient after the hemodilution was performed twice. In contrast, the COLs in the limb muscles and kidneys started to fluctuate and/or recover slowly even after the first hemodilution. After the fourth hemodilution, arterial hypotension developed and the brain COL decreased to approximately one-fifth of the initial value. During this period of cerebral ischemia, administering 1 μg of adrenaline for resuscitation resulted in a sudden augmentation of the systemic BP. Remarkably, the brain COL showed substantial recovery, but muscle and kidney COLs quickly decreased to nearly zero, supporting the brain-oriented vasodilation effects of this catecholamine compound in crisis^32–34^. Another shot of adrenaline (1 μg) ended its life.

### O_2_ measurement in the sanctuary

OxyBLI consists of a mutant of the firefly luciferase, which requires ATP in addition to O_2_ and the substrate for light emission^12^. The advantages and disadvantages of this ATP-dependent BLI system have been compared with those of the ATP-independent, coelenterazine-based BLI system^35,36^. The latter system is exemplified by a combination of the Nanoluc enzyme and its high-performance substrate, Furimazine^37^. Of particular importance, however, is the leakage of intracellularly expressed luciferases into interstitial fluids and circulating blood. When Furimazine was subcutaneously administered on the back of an Emx1-Nanoluc mouse (Fig. 10A), a strong bioluminescent signal was immediately observed at the injection site, increasing in intensity over a span of 50 min. Only negligible signals were detected in the brain, extremities, and kidneys. Furthermore, we collected blood samples from Emx1-Nanoluc mice for *in vitro* bioluminescence measurements (Fig. 10B). Remarkably, the addition of Furimazine increased the bioluminescence intensity by over three orders of magnitude, and this result was obtained reproducibly in five independent experiments using different mice. In contrast, subcutaneous administration of AkaLumine-HCl produced signals not at the injection site but in the brain, extremities, and kidneys of an Emx1-Venus-Akaluc mouse (Fig. 10C). It was also confirmed that no increase in bioluminescence intensity was detected in blood samples with the simple addition of AkaLumine. However, the further addition of ATP increased the intensity by approximately one order of magnitude (Fig. 10D). These results indicate that OxyBLI generates bioluminescent signals exclusively inside cells with abundant ATP. We thus argue that OxyBLI is suitable for O_2_ imaging in the sanctuary (Fig. 1, ocher-shaded area), all the better for its ATP dependency. In contrast, *in vivo* BLI systems using Nanoluc may be a sensitive tool for detecting dynamic changes in O_2_ levels in the intravascular space. Such systems could be useful for visualizing hemodynamics; for example, the localization of a fusion of Nanoluc and mNeonGreen, which is called GeNL^38^, in astrocytic networks enabled Beinlich et al. to discover and characterize activity-dependent hypoxic pockets in the cerebral cortex of awake mice^39^.

**Figure 10.**
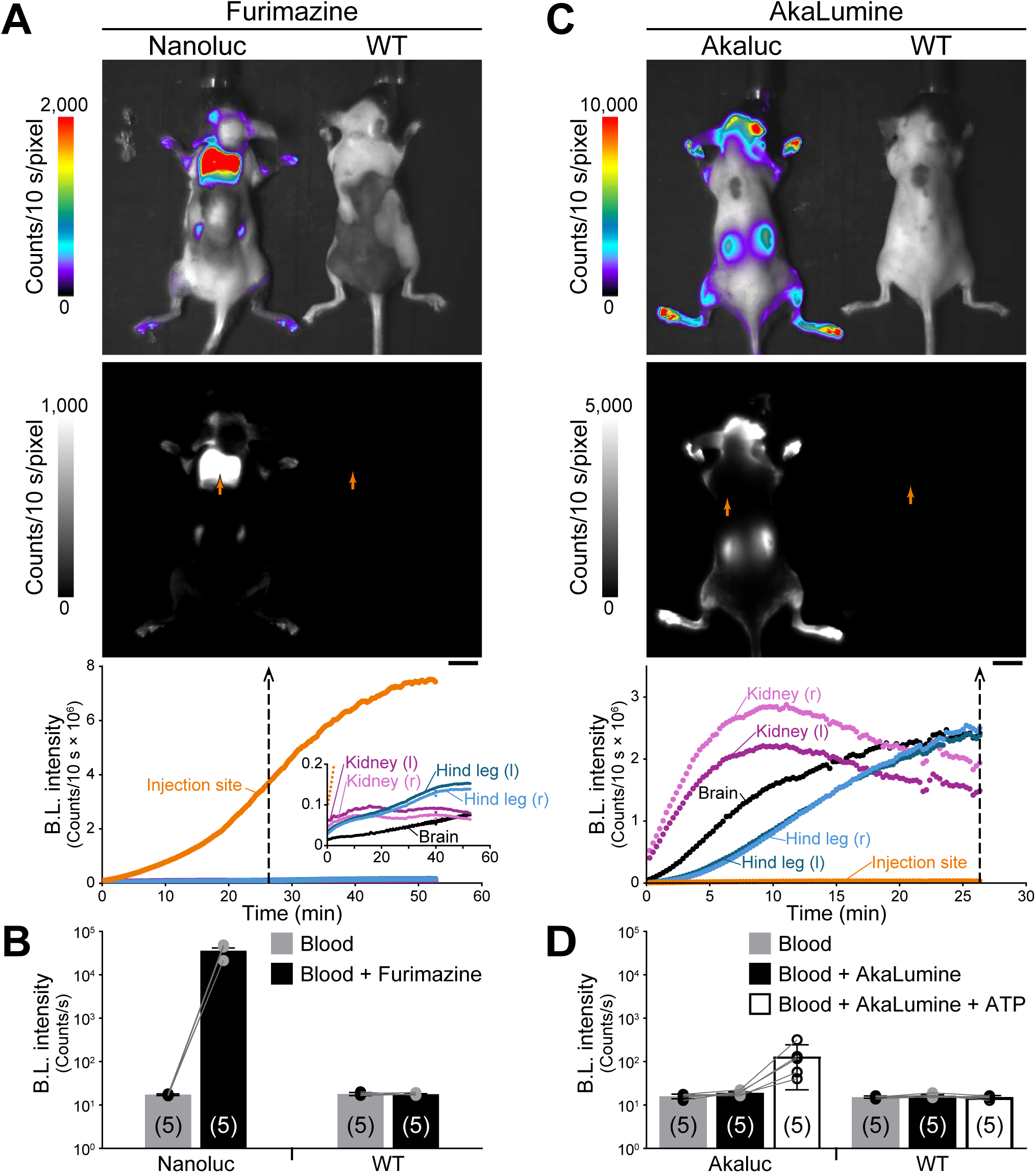
Comparison of ATP-dependent and ATP-independent BLI systems. **A,** *In vivo* BLI after subcutaneous administration of Furimazine on the back. An Emx1-Nanoluc mouse (left) and a wild-type (WT) mouse (right) were imaged simultaneously. Displayed in color (top) and black and white (bottom). The substrate injection sites are indicated by orange arrows in the black-and-white image. Scale bar: 10 mm. Shown below are the time courses of the bioluminescence signals in the injection site, brain, kidneys, and hind legs of the Emx1-Nanoluc mouse. *inset,* Development of signals from Nanoluc-expressing regions. **B,** *In vitro* bioluminescence measurements in the presence and absence of Furimazine. *left,* Emx1-Nanoluc mouse. *right,* wild-type (WT) mouse. Five blood samples from different mice were examined comparatively. Individual signal intensities are plotted as dots, and the mean values are shown as bars. **C,** *In vivo* BLI after subcutaneous administration of AkaLumine-HCl on the back. An Emx1-Venus-Akaluc mouse (left) and a wild-type (WT) mouse (right) were imaged simultaneously. Displayed in color (top) and black and white (bottom). The substrate injection sites are indicated by orange arrows in the black-and-white image. Scale bar: 10 mm. Shown below are the time courses of the bioluminescence signals in the injection site, brain, kidneys, and hind legs of the Emx1-Venus-Akaluc mouse. *inset,* Development of signals from Venus-Akaluc-expressing regions. **D,** *In vitro* bioluminescence measurements in the presence and absence of AkaLumine and/or ATP. *left,* Emx1-Venus-Akaluc mouse. *right,* wild-type (WT) mouse. Five blood samples from different mice were examined comparatively. Individual signal intensities are plotted as dots, and the mean values are shown as bars.

## Discussion

OxyBLI enables the direct visualization of tissue oxygenation dynamics in intact animals, offering a comprehensive understanding of the respiratory and circulatory mechanisms that regulate systemic O_2_ delivery. In this study, we confirmed the effects of common clinical interventions, including isovolemic hemodilution against bleeding, blood transfusions for anemia treatment, adrenaline administration for brain-directed oxygenation during cardiopulmonary resuscitation, and hypoventilation for improved brain tissue oxygenation via permissive hypercapnia. Here, however, we will focus our discussion mainly on findings useful for assessing how tissue oxygenation is affected by changes in FiO_2_, particularly hyperoxic breathing, as there are many uncertainties surrounding this topic. The advantages of OxyBLI over conventional electrode-based methods are detailed in Supplementary Discussion.

### O_2_-induced systemic vasoconstriction

In the absence of a lung disorder that impairs external respiration, PaO_2_ is almost linearly correlated with FiO_2_ across the full range. We confirmed this relationship for PaO_2_ values below 200 mmHg using an OxyLite sensor in the rat femoral artery (Fig. 4). We also quantified hyperoxemia in rats subjected to hyperoxic breathing (1.0 FiO_2_) using BGA. The average PaO₂ value was 424 ± 36 mmHg, which is indeed several times higher than the value at 0.21 FiO_2_ (Fig. 11). However, the relationship between PaO_2_ and tissue O_2_ level is largely nonlinear. The merged traces of the femoral arterial PaO_2_ and striatal COL, originating from Figure 8C, are illustrated in Figure 11. The two were well correlated between 0.21 and 0.1 FiO_2_, but not at 1.0 FiO_2_. Therefore, it is highly unlikely that hyperoxemia simply results in tissue hyperoxia. This is because O_2_ functions as a strong vasoconstrictor in all major organs except the lungs.

**Figure 11.**
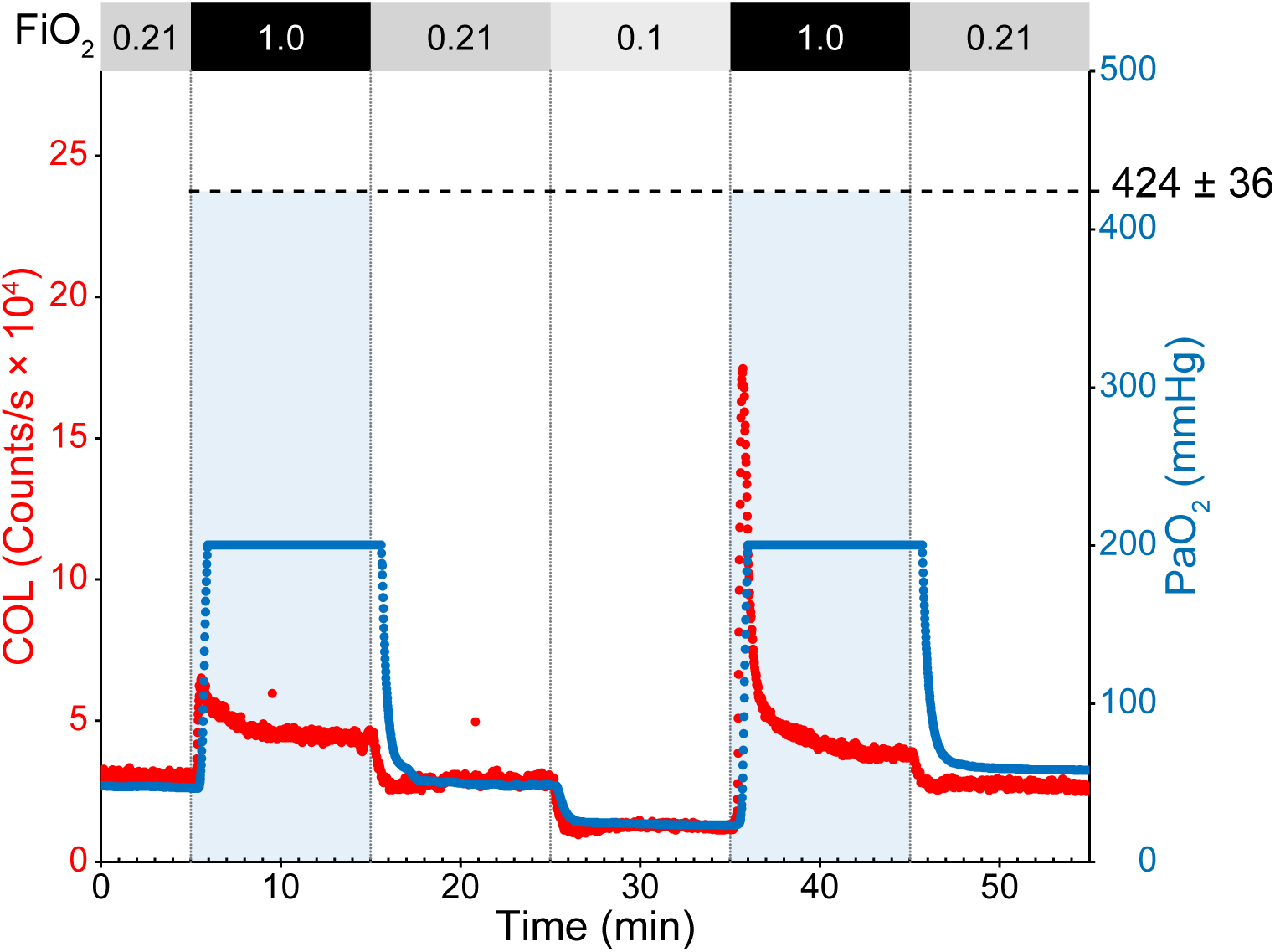
Relationship between striatal COL, PaO_2_ and FiO_2_. Time courses of striatal COL (red dots) and femoral arterial PaO_2_ (blue dots) during the regular FiO_2_ protocol experiment shown in Figure 8C. Thery were superimposed with their baselines at 0.21 FiO_2_ aligned. Due to the 5-second moving average of the OxyLite sampling, there was a slight delay in the PaO_2_ response to a change in FiO_2_. Although the rat had no underlying lung disease, the measured PaO_2_ values were slightly decreased, likely due to the effects of artificial ventilation. The PaO_2_/FiO_2_ ratio was approximately 350 mmHg. Due to the OxyLite sensor’s maximum detection limit of 200 mmHg, the PaO_2_ at 1.0 FiO_2_ was measured by BGA to be 424 ± 36 mmHg (*n* = 4 experiments). The average value was superimposed on the graph.

The vasoconstrictor effect of O_2_ is evidenced by decreased capillary density in *in situ* skeletal muscle preparations^40^. Also, *in vivo* systemic vasoconstriction induced by O_2_ has been demonstrated using the hamster window chamber model, which maintains tissue integrity well. Tsai et al. conducted an intravital microscopy study to examine the microvasculature of awake animals under normoxic and hyperoxic conditions^41^. After exposing the animals to 1.0 FiO_2_ for 30 min, they measured various experimental parameters and found that interstitial O_2_ tension increased by only 47% compared to the control group (0.21 FiO_2_), due to significant decreases in cardiac output, capillary flow, and capillary density. Through *in vivo* OxyBLI experiments using rats and mice, we found that brain COL gradually reached steady-state levels after overshooting under hyperoxic conditions. Steady-state levels in artificially ventilated rats at 1.0 FiO_2_ were, on average, 41.4% and 44.1% higher than the COL baseline at 0.21 FiO_2_ when FiO_2_ was increased from 0.21 and 0.1, respectively (Fig. 6B, right). This nearly 50% increase in COL supports the existence of such vasoconstriction regulation. By contrast, in our *ex vivo* OxyBLI experiments using the detached mouse auricle, where the microvasculature was decoupled from systemic regulation, no saturation (no vasoconstriction) was detected in the COL (Fig. 3; Supplementary Figs. 2 and 3). The increment of auricular COL at 1.0 FbO_2_ was significant with 1,166 ± 156% (*n* = 10 experiments using 6 Emx1-Venus-Akaluc mice) or 916 ± 154% (*n* = 6 experiments using 4 CAG-Venus-Akaluc mice) compared to that at 0.21 FbO_2_.

Although numerous studies have investigated tissue oxygenation in hypoxic and/or hyperoxic conditions by changing FiO_2_ settings, its dynamic aspects during the transition state have hardly been explored. In the aforementioned study by Tsai et al., for example, a 30-min adaptation period was set between the initiation of hyperoxic breathing (1.0 FiO_2_) and actual measurements^41^. However, such an experimental design would miss the overshooting responses that we observed, which lasted for no more than 5 min. Therefore, real-time observation of tissue oxygenation is crucial for understanding microvasculature dynamics.

### Tissue microvasculature for vasoconstriction

O_2_-induced vasoconstriction occurs at the transition from arterioles to capillaries. In physiology and histology textbooks, the general architecture of the microcirculation is illustrated with schematic diagrams that highlight precapillary sphincters and metarterioles (Fig. 12A)^42,43^. Although the existence of these two components in the brain microcirculation is not clear, we use the textbook diagrams to interpret our data on rat brain vasoconstrictor responses. In principle, hyperoxemia owing to 1.0 FiO_2_ causes considerable contraction of the precapillary sphincters, ultimately reducing blood flow into the capillary bed. This regulation limits the increase in COL until it reaches steady-state levels. On the other hand, metarteriolar regulation, if it exists, appears to be rather complex. The time courses of CBF were explicitly different when FiO_2_ increased to 1.0 from 0.21 vs. 0.1 (Figs. 4D, 5, and 8C). The 1.0 FiO_2_ challenge from normoxia (0.21 FiO_2_) decreased CBF slightly, suggesting an increase in precapillary resistance possibly due to precapillary sphincter contraction (Fig. 12B). Conversely, the 1.0 FiO_2_ challenge from hypoxia (0.1 FiO_2_) resulted in a transient but considerable increase in CBF. This FiO_2_ challenge reflects the situation in which patients with acute systemic hypoxia receive emergency O_2_ administration. The significant increase in CBF, which is more sustained than the COL overshoot, may be explained by an accelerated arteriovenous shunt through the metarterioles and the thoroughfare channels (Fig. 12C). Hypoxia-induced alterations in sensitivity to O_2_ may lead to the relaxation of metarterioles as well as the contraction of precapillary sphincters, thereby efficiently bypassing the capillary bed.

**Figure 12.**
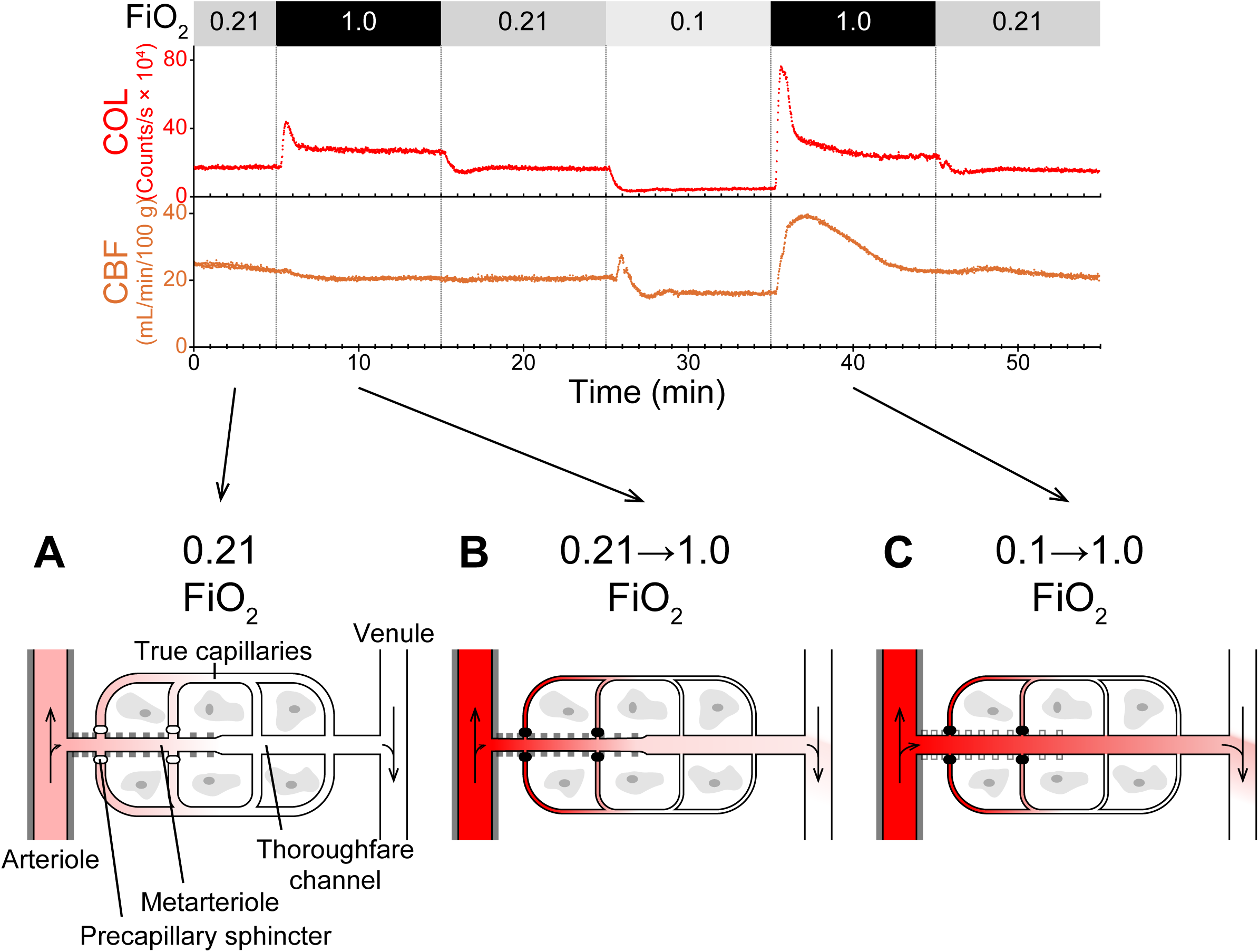
Schematic diagrams illustrating the microcirculation. The COL and CBF data from Figure 4D are shown above. Below are three assumed microcirculation situations: under normoxia (**A**), hyperoxia after normoxia (**B**), and hyperoxia after hypoxia (**C**). The basic architecture of the microcirculation is illustrated based on classic observations of mesenteric vasculature^42,43^. True capillaries branch from a metarteriole, which are enriched by scattered smooth muscle fibers (indicated by gray stripes), and converge into a thoroughfare channel that lacks muscle. Precapillary sphincters (indicated by ovals) regulate blood flow into the network of true capillaries. **A,** Normoxic condition. This diagram has annotations. The precapillary sphincters are open (open ovals). The metarteriole is slightly contracted. **B,** Hyperoxic condition after normoxia. This situation corresponds to the 1^st^ phase of the regular FiO_2_ protocol. Hyperoxemia (high PaO_2_, indicated by dark red) leads to the closure of the precapillary sphincters (solid ovals). As a result, precapillary resistance increases slightly, and CBF decreases accordingly. **C,** Hyperoxic condition after hypoxia. This situation corresponds to the 2^nd^ phase of the regular FiO_2_ protocol. In addition to hyperoxemia and precapillary sphincter closure, the metarteriole fully relaxes allowing for a more efficient bypass of the capillary beds, which may account for the paradoxical increase in CBF.

### Relationship between OxyBLI and AkaBLI

It should be noted that OxyBLI, which consists of AkaLumine and Akaluc, is essentially equivalent to AkaBLI. When its O_2_ dependency is focused on, it is called OxyBLI. Our present study does not intend to undermine the value of AkaBLI. With adequate consideration of FiO_2_, AkaBLI enables comprehensive, quantitative measurements of biological processes in living animals owing to the homogeneous biodistribution of the AkaLumine substrate. In all events, FiO_2_ is critical metadata. Every BLI system is more or less sensitive to O_2_ concentration. Notably, some classic *in vivo* BLI protocols involve a hyperoxic environment (1.0 FiO_2_) (refs. 44, 45), which should bring about the O_2_ regulations that we observed in this study.

### Limitations of the study

We will point out three limitations of OxyBLI. First, its readout (i.e., COL) does not provide absolute PO_2_ values, which are usually obtained by *in vitro*-calibratable, time-resolved, O_2_-quenching measurements. Remarkably, however, the OxyBLI baseline at 0.21 FiO_2_ remains considerably stable for hours; the COL signals were neither normalized nor corrected in this study. Additionally, our *ex vivo* experiments revealed the system’s broad linearity. These features of OxyBLI should enable quantitative measurements of changes in tissue PO_2_. Second, the bioluminescent readout of OxyBLI naturally has poor spatial resolution. However, its practical brightness enables the visualization of genetically defined small cell populations^12^. Third, the consumption of O_2_ by OxyBLI may result in weak signals, as is typically the case with the electrochemical electrode (the Clark electrode) method. When O_2_ supply is extremely low, it is essential to consider the amounts of O_2_ consumed by Akaluc, in addition to that consumed by cellular mitochondria, for the quantitative assessment of COL.

### Clinical perspectives

In this study, we subjected healthy mice and rats to conditions that corresponded to those of acutely ill patients. A simple question arises: Is routine O_2_ therapy really required for all patients admitted to the intensive care unit? Emergency physicians must avoid not only hypoxia but also hyperoxia. However, the extent of tissue hyperoxia associated with supplemental O_2_ and the resultant hyperoxemia remains unknown. Although the conventional threshold for O_2_ therapy and its target are set based on SpO_2_ in clinical medicine, arterial O_2_ saturation measured by pulse oximetry varies markedly from actual tissue oxygenation levels^8^. In principle, O_2_-induced vasoconstriction protects tissues from exposure to excess O_2_, and systemic vasoconstriction increases left ventricular afterload, resulting in decreased cardiac output. Overall, this type of response can explain why supplemental O_2_ has little to no influence on whole-body O_2_ consumption^8^.

The direct measurement of COL by OxyBLI enabled us to conclude that the increase in tissue PO_2_ during O_2_ administration (1.0 FiO_2_) is relatively mild in the steady state, only approximately 50% compared with the baseline level. It remains an open question, however, how this mild increase in PO_2_ affects tissue homeostasis. So, the risk of hyperoxic tissue injury must be assessed not only in the lungs but also in other organs^46^. During the transition period, on the other hand, severe hyperoxia occurs but only for a moment. Real-time OxyBLI measurements enabled us to discover and characterize the overshooting components of tissue PO_2_ immediately after the initiation of O_2_ administration. Because its peak amplitude depends on the degree of tissue hypoxia prior to the procedure (Figs. 2, 4, 5 and 6), this transient tissue hyperoxia may ameliorate the O_2_ deficiency in tissues. Otherwise, the overshoot may result in excessive O_2_ delivery, causing unwanted oxidative stress. Is overshooting tissue hyperoxia beneficial or harmful? We believe this issue is better left to a future study, once we or others have developed better experimental designs.

There also remains clinical uncertainty regarding the effectiveness of hyperoxia in preventing surgical site infection (SSI)^47,48^. Although the 2016 World Health Organization (WHO) guidelines recommend the perioperative administration of high-dose O_2_ (80%) for the purpose of reducing SSI, anesthesiologists are earnestly seeking the optimal concentration of O_2_ to be administered. Currently, 30% to 100% is a typical range. As a piece of valuable information, it is noted that a slightly high dose of 30–40% was sufficient to saturate COL to the steady state (Fig. 4B). The O_2_ concentrations for this saturation may be reminiscent of the highest atmospheric O_2_ (35%) in Earth’s history and may be used as the minimum effective doses. Future applications of OxyBLI in nonclinical studies of respiratory and circulatory physiology will provide a wealth of information on tissue oxygenation regulation. Such information will lay the foundation for the development of new markers for evaluating the risks and benefits of clinical interventions, including O_2_ administration, on critically ill patients.

## Data, code, and materials availability

All data generated in this study are available through the RIKEN Research Data and Copyrighted-work Management System (dmsgrdm.riken.jp/egdq4/). Materials are available from the lead contact, A.M. upon reasonable request.

## Acknowledgments

The authors thank the following people and organizations for their contributions to this study: Dr. Masahiro Jinzaki (Keio Univ.) for his research coordination; Mengtong Zhang, Jue Wang, and Yizhi Geng (Keio Univ.) for their technical assistance; the RIKEN CBS-EVIDENT Open Collaboration Center (BOCC) for their assistance in microscopy observation; Kurogane Kasei Co., Ltd. for their material support; Dr. Masamichi Yamamoto (The National Cerebral and Cardiovascular Center) for ATP measurement in biological samples; Dr. Kengo Inoue (Univ. of Miyazaki) for assistance with *in vitro* experiments under anaerobic conditions; Dr. Yuki Hiruta (Keio Univ.) for Furimazine provision; Dr. Hiroshi Kurokawa (RIKEN CBS) for assistance with statistical analysis; Dr. Shigeo Okabe (The Univ. of Tokyo), Dr. Tetsuya Matsuda (Tamagawa Univ.), Dr. Toshihisa Ohtsuka (Univ. of Yamanashi), and Dr. Masayuki Watanabe (Hokkaido Univ.), and Drs. Takashige Yamada and Hiroshi Morisaki (Keio Univ.) for their continuous support.

## Author contributions

S.I., J.K., and A.M. designed *in vivo* OxyBLI experiments using mice. S.I., J.K., and T.T. performed the mouse experiments and analyzed the data. J.K. designed *ex vivo* OxyBLI experiments using detached mouse auricles. J.K. and S.I performed the *ex vivo* OxyBLI experiments and analyzed the data. J.K., T.T., and A.M. designed *in vivo* OxyBLI experiments using rats. J.K. and T.T. performed the rat experiments and analyzed the data. H. Hama performed histological experiments. S.I., R.T., and T.N. prepared luciferase-expressing mouse lines. S.I. performed comparative BLI experiments using different luciferin/luciferase pairs. S.I. and M.S. quantified COL dynamics. H. Hioki and M.T. prepared AAVs, which H. Hama, M.S., and R.T. then injected into rat brains. A.M. and M.S. prepared the figures. J.K. integrated systemic parameter measurements with surgical procedures, and supervised the research from a clinical perspective. A.M. designed and wrote the manuscript. A.M. supervised the whole project.

## Additional information

Supplementary information accompanies this paper.

## Materials and Methods

### Animals

The experimental procedures and housing conditions for the animals were approved by the Animal Experiments Committee at RIKEN, Keio University School of Medicine, and University of Miyazaki. All animals were cared for and treated humanely in accordance with the institutional guidelines for experiments using animals. All mice and rats used for OxyBLI in this study are listed in Supplementary Tables 1 and 2, respectively. All mice used for immunohistochemistry are listed in Supplementary Table 3.

### AkaLumine

Due to the limited volume for intravenous or intraperitoneal administration (∼200 µL for a mouse and ∼500 µL for a rat), *in vivo* OxyBLI as well as AkaBLI require AkaLumine substrate concentrations higher than 1 mM^12^. Thus, AkaLumine is administered to animals in the form of its hydrochloride salt, AkaLumine hydrochloride (AkaLumine-HCl), which is highly water soluble. AkaLumine-HCl is commercially available under the name TokeOni from the following three companies:

Kurogane Kasei Co., Ltd. in Japan (https://www.kuroganekasei.com/en/development-tokeoni.html), Sigma-Aldrich Co., LLC, St. Louis, MO, USA (https://www.sigmaaldrich.com/US/en/product/aldrich/808350), Summit Pharmaceuticals International Corporation, in Japan (https://www.summitpharma.co.jp/japanese/service/products/reagents_lumi/ivi.html). AkaLumine-HCl is also purchasable from FUJIFILM Wako Pure Chemical Corporation (https://labchem-wako.fujifilm.com/us/product/detail/W01W0101-2670.html).

In the present study, a stock solution of 60 mM AkaLumine-HCl (TokeOni, Kurogane Kasei Co., Ltd., Japan) was stored in aliquots at −80°C.

### AAV infection in the mouse striatum

An ICR mice (male, 10 weeks old; Japan SLC, Inc.) was anesthetized with pentobarbital (65 mg/kg BW) and mounted onto a stereotaxic frame. A 0.5-µL solution of AAV2/1 SynTetOff Venus-Akaluc (2.0 × 10¹⁰ IFU/mL) was injected into the striatum using a micro pump (Muromachi Kikai Co., Ltd., Tokyo, Japan). The injection site was 0.14 mm anterior to the bregma, 2.1 mm lateral to the midline, and 2.8 mm deep from the brain surface. pAAV2 SynTeOff Venus-Akaluc was used to produce the recombinant AAV (AAV2/1 SynTetOff Venus-Akaluc) vectors as described previously^13^. BLI was performed forty weeks after the AAV injection. See Fig. 2A.

### Mouse striatal OxyBLI

An ICR mouse expressing Akaluc in the striatum was imaged in a prone position under inhalation anesthesia with 2% isoflurane (isoflurane for animals, MSD Animal health, Japan) via a nose cone after intraperitoneal administration of 100 µL of 30 mM AkaLumine-HCl (75 nmol/g BW). BLI was performed using a Lumazone system (Nippon Roper, Tokyo, Japan) equipped with an iKon L camera (Andor Technology Ltd., UK) and an AI Nikkor lens (φ50 mm, f: 1.2S; Nikon Corporation, Japan). The dorsal surface was imaged continuously at a rate of 0.1 frames per second with 16×16 binning and a gain of 4. A bright-field image was acquired for 30 msec at the end of the experiment. Image acquisition was carried out using Andor Solis software (ver. 4.29, Andor Technology Ltd., UK), and image analysis was performed using MetaMorph software (ver. 7.10, Molecular Devices, LLC., Sunnyvale, CA, USA).

Hypoxic breathing was achieved by adding nitrogen (N₂) gas to air via a regulator. The O_2_ partial pressure emerging from the nose cone was measured in advance using an Oxyman (OM-25MF10, Taiei Engineering, Japan) O_2_ monitor device to determine the required air:N₂ mixture ratio to obtain 0.1 and 0.05 FiO₂. See Fig. 2A.

### Mouse preparation for *in vivo* OxyBLI during spontaneous breathing

CAG-loxP-STOP-loxP (LSL)-Venus-Akaluc mice (RBRC10858, RIKEN BRC, Japan) were crossbred with the following Cre drivers: Emx1-Cre (RBRC01345, RIKEN BRC, Japan); VGAT-Cre (RBRC10606, RIKEN BRC, Japan); αMHC-Cre (https://pmc.ncbi.nlm.nih.gov/articles/PMC533965/); and Alb-Cre (003574, The Jackson Laboratory, USA). The following mouse lines were generated: Emx1-Venus-Akaluc, VGAT-Venus-Akaluc, αMHC-Venus-Akaluc, and Alb-Venus-Akaluc. After anesthetizing the mouse with 2% isoflurane in an 18 × 18 × 20 cm anesthesia chamber (SN-487-85-02, Shinano Manufacturing Co., Ltd., Japan), approximately 80 µL of 30 mM AkaLumine-HCl (75 nmol/g BW) was administered intraperitoneally or intravenously. Thirty to eighty minutes after AkaLumine-HCl administration, BLI was performed while the mouse received 2% isoflurane via a nose cone. Hypoxic breathing was achieved using a 10% O2 cylinder (composition: O₂ 10%, N₂ 88.96%, CO₂ 0.04%). Hyperoxic breathing (1.0 FiO2) was achieved using a 100% O2 cylinder. The following BLI systems were used:

A MIIS system (Molecular Devices Japan, Tokyo, Japan) equipped with an Evolve 512 EMCCD camera (Photometrics, Tucson, AZ, USA) and a YMV1795 lens (φ17 mm, f: 0.95; Yakumo Optical Corp.) was used in the experiments depicted in Figure 2B. A home-made system equipped with the same camera and lens was used in the experiment depicted in Figure 4A.

### Mouse preparation for *in vivo* OxyBLI during mechanical ventilation

Emx1-Venus-Akaluc mice were first administered approximately 40 µL of 60 mM AkaLumine-HCl (75 nmol/g BW) intraperitoneally while awake. Then, anesthesia was induced using 3% isoflurane (Wako, 099-06571) in a 10 ×10 × 10 cm anesthesia chamber (anesthesia machine: SN-487-OT, Shinano Manufacturing Co., Ltd.).

For tracheotomy and endotracheal intubation, the mouse was placed in the supine position on a heated monitoring platform (Small Animal Physiological Monitoring System, Harvard Apparatus), which was maintained at 37°C. Anesthesia was maintained with 2–2.5% isoflurane/air administered via a nose cone. Butorphanol (Vetorphale®, Meiji Animal Health) (2.5 mg/kg) was administered intraperitoneally for analgesia. After shaving the neck hair, a 1-cm midline incision was made in the neck. Right-angled forceps were used for blunt dissection. The exposed trachea was then incised at the thyroid level using the tip of a 23-gauge needle. A 22-gauge venous catheter (Surflon 22G, outer diameter 0.85 mm, length 25 mm) was inserted approximately 5 mm into the trachea. The insertion site was secured with 5-0 braided silk thread (Shirakawa, DMB501002). Mechanical ventilation was performed at a tidal volume of 8 mL/kg and a respiratory rate of 70/min using a mouse ventilator (Minivent type 845, Harvard Apparatus), which was connected to the catheter. An electrocardiogram (ECG) was used for vital monitoring, and an SpO₂ probe (Small Animal Physiological Monitoring System 75-1500, Harvard Apparatus) was placed on the tail.

While in a prone position, the hair on the head was shaved and the skin was exposed to depilatory cream (Epilat, Kracie). Lastly, the mouse was moved to a stage inside a dark box (80 cm × 80 cm × 90 cm), which was enclosed by an opaque black curtain, for OxyBLI.

See Figs. 3A, 3B and 9B.

### Hindlimb compression model

A Microcuff ET tube (I.D. 3.0 mm; Avanos Medical, Inc.) with a cuff was pressed against the ventral aspect of the mouse’s right thigh and secured with 1.2 cm wide, double-sided, hook-and-loop fastener tape (Easy-Wrapping Belt, NC2232; 3M Japan Ltd., Tokyo, Japan). During leg compression, the cuff was inflated with 2.5 mL of air. To monitor distal blood flow, a laser Doppler flow meter (ALF21, ADVANCE Co., Ltd., Tokyo, Japan) was pressed against the heel of the right hind paw and secured with clay. See Fig. 9A.

### Isovolumic hemodilution model

The right carotid artery of the mouse was bluntly dissected for cannulation. The distal end of the cannulation site near the external carotid bifurcation was ligated with 5-0 silk thread, while the proximal end was clamped with a 7-mm curved vascular clip. A 26-gauge venous catheter (Supercath 5, outer diameter 0.6 mm, length 19 mm; Togo Medikit Co., Ltd.) was inserted and secured with 5-0 silk thread. The other end of the catheter was connected to a polyethylene tube (PE-50, EastsideMed). The vascular clip was then removed. After confirming arterial blood drainage, the surgical site was flushed with 0.5 mL of a saline solution containing two units of heparin (Fuso Pharmaceutical Industries, Ltd.). The polyethylene tube was connected to an arterial blood pressure monitor (Small Animal Physiological Monitoring System, Harvard Apparatus).

For isovolumic hemodilution, 0.5 mL of blood was withdrawn from the arterial line over approximately 30 seconds, followed by the injection of 0.5 mL of saline solution into the arterial line over a similar time period. Adrenaline (Bosmin®, Daiichi Sankyo Company Ltd.) was diluted in saline to 10 µg/mL. One microgram was injected via the arterial line, immediately followed by flushing the arterial line with 0.5 mL of saline.

See Fig. 9B.

### AAV infection in the rat striatum

Five-week-old male Sprague-Dawley rats (Japan SLC, Inc.) were anesthetized using 2% isoflurane and a combination of intraperitoneally administered medetomidine (0.3 mg/kg BW), midazolam (2 mg/kg BW), and butorphanol (5 mg/kg BW), and mounted onto a stereotaxic frame (Model 900, David Kopf Instruments). Stereotaxic surgery was performed bilaterally (coordinates: AP = 0.96, ML = ±3.0, DV = 4.16). After administering local anesthesia with 8% Xylocaine (Sandoz) and disinfecting the areas with Povidone-Iodine (Meiji Seika Pharma), incisions were made in the scalp. Then, the skull was exposed and the periosteum was detached. Lastly, a 1-mm-diameter hole was drilled into the skull on each side using a dental drill (Viva Mate G5, NSK, Japan). A 1.2-µL solution of AAV2/1 SynTetOff Venus-Akaluc (1.02 × 10^14^ gc/mL) was injected into the right striatum using a micro pump (Model 310 Plus, KD Scientific). BLI was performed at least two weeks after the AAV injection. The hole on the left was used to insert a laser Doppler flowmeter probe. See Figs. 4, 5, 7 and 8.

### Rat preparation for *in vivo* OxyBLI during mechanical ventilation

Rats with striatal expression of Venus-Akaluc were used. Anesthesia was induced with 3% isoflurane in an anesthesia chamber in the same manner as for mice. The rats were then placed in a supine position on a 37°C surgical table. For subsequent procedures, anesthesia was maintained with 2% isoflurane/air delivered via a nose cone. Analgesia was induced via intraperitoneal administration of butorphanol (2.5 mg/kg). At this time, 400 µL of 60 mM AkaLumine-HCl was administered intraperitoneally.

After shaving the hair, a skin incision was made in the left inguinal region and the femoral artery was bluntly dissected for cannulation. The distal end of the cannulation site was ligated with 5-0 silk thread while the proximal end was clamped with a 7-mm curved vascular clip.

Then, a 24-gauge venous catheter (Supercath 5, O.D. 0.7 mm, length 19 mm; Togo Medikit Co., Ltd.) was inserted and secured with 5-0 silk thread. The other end of the catheter was connected to a polyethylene tube (PE-50, EastsideMed). Then, the vascular clip was removed. After confirming arterial blood drainage, the surgical site was flushed with 0.5 mL of saline solution containing 2 units of heparin (Fuso Pharmaceutical Industries, Ltd.). The polyethylene tube was then connected to an arterial blood pressure monitor (Small Animal Physiological Monitoring System, Harvard Apparatus). The right femoral artery was exposed in the same way for cannulation. A 24-gauge venous catheter was used to place an OxyLite™ probe (tip diameter: NX-BF/OT/E with a tip diameter of 350 µm, a luminescence wavelength of 650 nm, and a data output of 1 Hz) into the femoral artery. The tip was positioned 5 mm away from the insertion site. To eliminate spontaneous breathing, pancuronium (Mioblock®, 4 mg; Schering-Plough) was administered intraperitoneally.

For the tracheotomy, a midline skin incision was made on the neck to expose the trachea. At the thyroid level, a 16-gauge venous cannula (outer diameter [O.D.] 1.7 mm, length 64 mm; Surflo, Terumo) was inserted approximately 1 cm and secured with 5-0 silk thread. The cannula was connected to a ventilator (SN-480-7, Shinano Manufacturing Co., Ltd.) and mechanical ventilation was performed at a rate of 8 mL/kg per minute.

The rat was then repositioned prone and an incision was made in the scalp. Through the hole previously created in the contralateral skull during AAV injection, a laser Doppler flowmeter probe (ALF21, ADVANCE; semiconductor laser, 780 nm, 10 Hz; BF04436, outer diameter [O.D.] 0.3 mm) was inserted approximately 5 mm into the brain toward the striatum. The probe was then secured to the skull with adhesive (Aron Alpha Extra, Konishi). For vital monitoring, an ECG was attached and an SpO₂ probe (Small Animal Physiological Monitoring System 75-1500, Harvard Apparatus) was placed on the right forepaw. Lastly, the rat was moved to a stage inside a dark box (80 cm × 80 cm × 90 cm), which was enclosed by an opaque black curtain, for OxyBLI.

See Figs. 4, 5, 7 and 8.

### Arterial blood gas analysis

Approximately 200 seconds after the onset of hyperventilation or hypoventilation, 0.2 mL of blood was taken from the right femoral artery of the rat for the measurement of pH, hemoglobin level (g/dL), PaO₂ (mmHg) and PaCO₂ (mmHg) using an i-Stat cartridge (Abbott, EG6+). See Figs. 7 and 11.

### Bleeding model

A volume of 2.5 mL was manually drained from the arterial line placed in the rat’s right femoral artery over approximately 30 seconds. Blood return was performed slowly over approximately one minute via the arterial line. For isovolumic hemodilution, 2.5 mL of blood was withdrawn from the arterial line over approximately 30 seconds and replaced with 2.5 mL of physiological saline over a similar time period.

Adrenaline (Bosmin®, Daiichi Sankyo Company Ltd.) was diluted in saline to 10 µg/mL in saline. Ten μg was injected via the arterial line, immediately followed by flushing the arterial line with 0.5 mL of saline. Cardiac arrest was induced by injecting 4 mmol of diluted KCl (Maruishi Pharmaceutical Co., Ltd.) through the arterial line. This was followed immediately by flushing the arterial line with 0.5 mL of physiological saline.

See Fig. 8.

### *In vivo* mouse OxyBLI

The experiments depicted in Figure 2B used a MIIS system (Molecular Devices Japan, Tokyo, Japan) equipped with an Evolve 512 EMCCD camera (Photometrics, Tucson, AZ, USA) and a YMV1795 lens (φ17 mm, f: 0.95, Yakumo Optical Corp.). The dorsal or ventral surface of the mouse was imaged continuously at a rate of 1–10 frames per second with no binning and a gain of 50–600. A bright-field image was acquired for 30 msec at the end of each experiment. Image acquisition was carried out using MetaMorph software (ver. 7.10, Molecular Devices, LLC., USA). Hypoxia (0.1 FiO₂) and hyperoxia (1.0 FiO₂) were achieved with 10% and 100% O_2_ cylinders, respectively. The composition of the 10% cylinder was as follows: O₂ (10%), N₂ (88.96%), and CO₂ (0.04%). The cylinders were switched using a regulator.

### *In vivo* mouse OxyBLI

The experiments depicted in Figures 3A, 3B and 9 used a home-made BLI system equipped with an Evolve 512 EMCCD camera (Photometrics, Tucson, AZ, USA) and a YMV1795 lens (φ17 mm, f: 0.95, Yakumo Optical Corp.). The dorsal surface of the mouse was imaged continuously at a rate of 1 frame per second with 2×2 binning and a gain of 100. Image acquisition was carried out using VisView (ver. 3.3.0.0, Visitron Systems). An FiO_2_ value of 0.1 was achieved by adding 100% N_2_ gas to air. Other gases with different FiO_2_ values were obtained using a combination of 100% O₂ and 100% N₂ cylinders.

### *In vivo* rat OxyBLI

The AAV-based TET-inducible system was used for the neuronal expression of Venus-Akaluc in the right striatum. Two to four weeks after the viral infection, rats were ip administered AkaLumine-HCl and subjected to OxyBLI under isoflurane anesthesia.

BLI was performed using a home-made BLI system equipped with an Evolve 512 EMCCD camera (Photometrics, Tucson, AZ, USA) and a YMV1795 lens (φ17 mm, f: 0.95; Yakumo Optical Corp.). The head was imaged continuously at a rate of 1 frame per second with 2×2 binning and a gain of 100. Two short-pass filters with cut-off wavelengths of 750 and 725 nm: SP750 and SP725 (#84-728 and #86-111, respectively; Edmund optics Japan, Japan) were used to eliminate the 780-nm light from the laser Doppler flow cytometer used for CBF measurement. Two SP750 filters and one SP725 filter were placed in front of the lens. These stacked filters pass most of the AkaLumine emission (Fig. 4A). Image acquisition was carried out using VisView (ver. 3.3.0.0, Visitron Systems).

Two hypoxic conditions (FiO₂ of 0.1 and 0.15) were achieved using 10% and 15% O_2_ cylinders with O₂:N₂ ratios of 10:90 and 15:85, respectively. Hyperoxic conditions (FiO₂ > 0.21) were created using a Dräger anesthesia machine with a mixture of air and 100% O₂.

Blood pressure (BP), oxygen saturation (SpO_2_), and cerebral blood flow (CBF) were sampled at frequencies of 250 Hz, 1 Hz and 10 Hz, respectively. Heart rate (HR) was monitored at a sampling frequency of 1 Hz from the ECG waveform. Although the OxyLite sampling rate was 1 Hz, the 5-second moving average slowed the response down. Analogue data were converted to digital format using a PowerLab device (AD Instruments) and recorded using LabChart software (also from AD Instruments). The data were then output in Excel format.

See Figs. 4, 5, 6, 7 and 8.

### *Ex vivo* mouse auricular OxyBLI

When using Emx1-Venus-Akaluc mice, *ex vivo* BLI experiments using the auricle were typically conducted after *in vivo* BLI experiments. Under 2–2.5% isoflurane anesthesia, the mouse auricle was excised at its base and mounted on clay (Blu Tack, Bostik Findley Pty. Ltd., Australia). The same gases used in the *in vivo* BLI experiments (FiO₂ of 0.1, 0.21, and 1.0) were blown at a rate of 1 L/min from a distance of approximately 5 mm using the outlet of a 5 mL syringe. The same *ex vivo* BLI experiments were performed on CAG-Venus-Akaluc mice, generated by crossing CAG-LSL-Venus-Akaluc with a CAG-Cre driver (Extended Data Fig. 3b, 4). BLI was performed using a home-made system equipped with an Evolve 512 EMCCD camera (Photometrics, Tucson, AZ, USA) and a TEC-M55 lens (φ55 mm, f: 2.8; CBC Co., Ltd.). The auricle was imaged continuously at a rate of 1 frame per second with 2×2 binning and a gain of 100. Image acquisition was carried out using VisView (ver. 3.3.0.0, Visitron Systems). See Figs. 3C and 3D; Supplementary Figs. 2 and 3.

### *Ex vivo* mouse auricular OxyBLI for dose-response relationships of firefly luciferase (FLuc) versus Akaluc

CAG-Venus-Akaluc and CAG-ffLuc-cp156 mice were used for comparison (https://www.sciencedirect.com/science/article/pii/S0006291X12001957). Under anesthesia with 2% isoflurane, they were administered respective substrates (AkaLumine-HCl, 75 nmol/g BW, and D-luciferin potassium salt, 500 nmol/g BW, In Vivo Grade, Promega, Madison, WI, USA). One to two hours after substrate administration, the auricles were detached for *ex vivo* BLI experiments. An FbO_2_ value of 0.1 was achieved by adding 100% N_2_ gas to air. Other gases with different FbO_2_ values were obtained using a combination of 100% O₂ and 100% N₂ cylinders. The O_2_ partial pressure emerging from the tube was checked in advance using a Microx4 Trace (PreSens Precision Sensing GmbH, Germany).

BLI was performed using a Lumazone system (Nippon Roper, Tokyo, Japan) equipped with an iKon L camera (Andor Technology Ltd., UK) and a lens (φ85 mm, f: 1.2, SPEEDMASTER for F mount, SHENYANG ZHONGYI OPTICAL TECHNOLOGY Co., Ltd., China). The ventral and dorsal surfaces were imaged continuously at 1–2 Hz with 8×8 binning and a gain of 4.

See Supplementary Fig. 3.

### Immunohistochemistry

Mice were deeply anesthetized using 2% isoflurane and a combination of intraperitoneally administered medetomidine (1.2 mg/kg BW), midazolam (12 mg/kg BW), and butorphanol (15 mg/kg BW), and then transcardially perfused with 4% (w/v) PFA/PBS(–). Brains, skeletal muscles, hearts, livers, kidneys, and auricles were removed and incubated in 4% (w/v) PFA/PBS(–) for post-fixation at 4°C for 1 d and then in 20% (wt/vol) sucrose/PBS(–) for cryoprotection at 4°C for 1 d. These samples were embedded in OCT compound (Tissue-Tek, Sakura) by using isopentane and liquid N_2_. Cryo-sections (35 μm thick) were prepared using a cryostat (CM3050S, Leica). The sections were permeabilized with 0.1% (w/v) Triton X-100 in PBS(–) at room temperature for 20 min. Then they were blocked with 2% (w/v) BSA in PBS(–) for1 h at room temperature. Lastly, they were processed using free-floating immunohistochemistry.

The following primary polyclonal antibodies (pAbs) and monoclonal antibodies (mAbs) were used in a solution of 0.1%(w/v) Triton X-100 in PBS(–).

Rat mAb to CD31 (1:100–300; BD Pharmingen, 553370) Rabbit pAb to GFP (1:400; Medical Biological Laboratory, 598) Goat pAb to desmin (1:300; LSBio, LS-B4463)

Sheep pAb to parvalbumin alpha (1:400; R&D Systems, AF5058) Mouse mAb to troponin T (1C11) (1:400; Abcam, ab8295) Secondary antibodies

The following secondary antibodies (Thermo Fisher Scientific) were used in 0.1%(w/v)

Triton X-100/PBS(–).

Goat anti-Mouse IgG (H+L) F(ab’)2 fragment, Alexa Fluor 633 conjugate (1:500; A21053) Goat anti-Rabbit IgG (H+L), DyLight 550 conjugate (1:1,000; SA5-10033)

Donkey anti-Goat IgG (H+L), Alexa Fluor 633 conjugate (1:500; A21082) Donkey anti-Sheep IgG (H+L), Alexa Fluor 633 conjugate (1:500; A21200) Goat anti-Rat IgG (H+L), Alexa Fluor 568 conjugate (1:500; A11077)

Additionally, the nuclei in skeletal muscles were directly labeled with SYTO61 (1:4,000; Thermo Fisher Scientific, S11343).

After each antibody reaction, the sections were rinsed three times with 0.1% (w/v) Triton X-100/PBS(–) at room temperature. Finally, the sections were equilibrated with 50% (w/v) glycerol in PBS(–) for 20 min at room temperature and mounted between slide and cover glasses. Samples were imaged using an upright laser scanning confocal microscopy system (Olympus FV1200) equipped with an XL Fluor 4×/340 (NA 0.28) (Evident) or U Plan Apo20× (NA 0.70) objective lens (Evident).

See Supplementary Figs. 1 and 2.

### Preparing mice for examining biodistribution of luciferase enzymes

Emx1-Naoluc mice were generated by crossbreeding CAG-DIO-Nanoluc mice with the Emx1 Cre driver. The CAG-DIO-NanoLuc mice with Cre-dependent Nanoluc expression were generated by the CRISPR/Cas9 technique using a targeting vector described in our previous publication^14^ with slight modifications. Briefly, synthetic DNA oligonucleotides for a DIO cassette consisting of two copies of loxP and lox2272 were generated and cloned between the CAG promoter and WPRE. A DNA fragment containing a NanoLuc coding sequence, which was custom-synthesized according to the nucleotide sequence information of NanoLuc (Promega, Madison, WI, USA), and an FRT site (Flp recombinase recognition site) was inserted within the DIO cassette. The resulting targeting vector (10 ng/μL) was injected into the pronuclei of zygotes derived from C57BL/6NJcl mice together with Cas9 protein (100 ng/μL; Thermo Fisher Scientific, Waltham, MA, USA), R26-T2 crRNA (50 ng/μL), tracrRNA (100 ng/μL), and Cas9 mRNA (25 ng/μL); these RNAs were generated by *in vitro* transcription using the T7-NLS hCas9-pA plasmid (RDB13130; RIKEN BRC, Japan). The crRNA and tracrRNA were purchased from FASMAC (Atsugi, Japan). The crRNA sequence was as follows: R26-T2-crRNA (5′-CGC CCA UCU UCU AGA AAG ACG UUU UAG AGC UAU GCU GUU UUG-3′). Genotyping primers are listed in our previous publication.

The Emx1-Venus-Akaluc mice were generated as described earlier^13^. All wild-type mice were C57BL/6J (Japan SLC, Inc.).

See Fig. 10.

### *In vivo* mouse BLI after subcutaneous administration of substrates

The entire body was depilated using clippers and depilatory cream (Epilat, Kracie Holdings, Ltd., Tokyo, Japan) while the mice were anesthetized with 2% isoflurane (Isoflurane for Animals, MSD Animal Health, Japan). While still under anesthesia, the mice were placed in a Lumazone dark box (Nippon Roper, Tokyo, Japan) on their stomachs, and the substrate was injected subcutaneously into their dorsal skin.

Furimazine, a gift from Dr. Hiruta at Keio University, was dissolved in a solution containing 35% (v/v) PEG-300 (164-09055, Fujifilm Wako Pure Chemical Corporation, Japan), 10% (v/v) glycerol (17017-35, Nacalai Tesque, Inc., Japan), 10% (v/v) ethanol (14722-75, Nacalai Tesque, Inc., Japan), and 10% (w/v) hydroxypropyl cyclodextrin (H0979, Tokyo Chemical Industry Co., Ltd., Japan). The final concentration of Furimazine was 2.73 mM. On the other hand, a solution of 30 mM AkaLumine-HCl was used. One hundred microliters of each solution were injected within 30 seconds.

Immediately after administration, BLI was performed using a Lumazone system (Nippon Roper, Tokyo, Japan) equipped with an iKon M camera (Andor Technology Ltd., UK) and a lens (M-5095C, φ50 mm, f: 0.95, M-5095C; Yakumo Optical Corp.). The mouse dorsal surfaces were imaged with 2×2 binning continuously at an exposure time of 10 sec for about 1 hour. Image acquisition was carried out using Andor Solis software (ver. 4.29, Andor Technology Ltd., UK), and image analysis was performed using MetaMorph software (ver. 7.10, Molecular Devices, LLC., Sunnyvale, CA, USA).

See Figs. 10A and 10C.

### *In vitro* bioluminescence measurement using blood samples

Under 2% isoflurane anesthesia, the mouse tail vein was incised with a scalpel. Emerging blood was collected and diluted tenfold with PBS containing heparin (1 U/mL). Samples from Emx1-Nanoluc, Emx1-Venus-Akaluc, and wild-type mice were refrigerated for up to two days.

The bioluminescence of each sample (19 µL) was measured in a test tube (55.476, Sarstedt AG & Co. KG, Germany) using an ATTO luminometer (AB-2280). First, measurements were performed using samples from five Emx1-Nanoluc mice and five wild-type mice. Then, after adding 1 µL of Nano-Glo Live Cell Reagent (N2011, Promega) containing Furimazine, the measurements were repeated. Similarly, measurements were performed using samples from five Emx1-Venus-Akaluc mice and five wild type mice, both before and after adding 1 µL of 2 mM AkaLumine-HCl, followed by an additional 1 µL of 21 mM ATP-Magnesium (00386-54, Nacalai Tesque, Inc.).

See Figs. 10B and 10D.

### Brain PO_2_ measurement using an optical electrode

An OxyLite probe (NX-BF/OT/E, Oxford Optronix, UK) was used to measure PO₂ inside the rat striatum. The probe has a 0.35 mm tip diameter, a 650 nm luminescence wavelength, and a 1 Hz data output rate. Measurements were repeated several times over the course of about one month.

A sheath was made from a 24-gauge intravenous catheter; the distal end was cut off at 5 mm, and the proximal end was slightly flared with heat. This sheath was used to guide the insertion of the OxyLite probe. Additionally, during implantation surgery and subsequent intervals, a 7-mm stylet made of 0.32-mm silver-plated copper wire (Artistic Wire®, Pakistan) was substituted for the probe.

On day 0, brain surgery was performed to place the sheath on a wild-type rat. Anesthesia was induced with 3% isoflurane and maintained with 2% isoflurane delivered via a nose cone. Additionally, analgesia was induced via intraperitoneal administration of butorphanol (2.5 mg/kg, Vetorphale®, Meiji Animal Health Co., Japan). After making a 10-mm incision in the scalp, a 1-mm burr hole was drilled in the skull at 1.0 mm anterior to the bregma and 3.0 mm lateral to the midline. After achieving hemostasis, the sheath with the stylet was advanced through the hole to a depth of 5 mm. In this position, the tip of the stylet was 7 mm deep from the brain surface, which was assumed to be in the middle of the striatum. The incision was closed with 6-0 nylon sutures (NA106, Crownjun, Japan), and bacitracin-fradiomycin sulfate ointment (Baramycin Ointment®, Toyo Seiyaku Kasei, Japan) was applied to the wound.

Prior to PO_2_ measurement, anesthesia and analgesia were administered as described above. After placing the rat in the prone position, the stylet was carefully replaced with the OxyLite probe through the sheath. The probe’s tip was confirmed to be 2 mm beyond the tip of the sheath. Representative PO₂ data obtained on days 6 and 23 are shown in Supplementary Figures 4B and 4C, respectively.

Electrocardiogram (ECG) electrodes were attached, and an SpO₂ probe (Small Animal Physiological Monitoring System, Model 75-1500, Harvard Apparatus) was placed on the right forepaw. Brain PO₂ data obtained using the OxyLite™ system were sampled at 1 Hz and processed using a five-second moving average. Analog signals were converted to a digital format using a PowerLab data acquisition system (ADInstruments) and recorded using LabChart software (ADInstruments). A premixed gas cylinder containing 10% O₂ and 90% N₂ was used to achieve the hypoxic condition (0.10 FiO₂), and 100% O₂ supplied from a cylinder connected to a Dräger anesthesia machine was used to achieve the hyperoxic condition (1.0 FiO₂).

### Measurement of hyperoxemia in rats under hyperoxic breathing

Male Sprague-Dawley rats were anesthetized and intubated in the same manner as in the experiments depicted in Figures 4, 5, 6, 7 and 8. The rats were exposed to hyperoxia (1.0 FiO₂), and blood gas analysis was performed on arterial blood samples drawn from the femoral arteries using an i-Stat cartridge (Abbott, EG6+). This experiment was repeated in three experiments using different rats.

### OxyBLI Analysis

OxyBLI data were analyzed using ImageJ version 2.3.0/1.54i or MetaMorph software Version 7.10. The data were opened as 16-bit files and ROIs were placed manually. The integrated density (total intensity) within each ROI was then measured every second. Background intensity was subtracted by measuring an ROI of identical area and shape within the same field of view, yielding the COL (counts per second per pixel).

## Supplementary information

### Supplementary Discussion

#### Advantages of OxyBLI over electrode-based methods

Aiming for the direct measurement of O_2_ levels in tissues, conventional methods use electrochemical or optical electrodes to determine O_2_ content in biological fluids in the extravascular space^10^. Compared with these electrode-based methods, our OxyBLI method has three advantages.

First, OxyBLI is a noninvasive method for monitoring COLs in intact cells. In contrast, conventional methods are invasive; the mechanical insertion of electrodes into tissues likely disrupts cells and small blood vessels, leading to false signals. Even with their chronic implantation, it is unclear to what extent the regulation of the microcirculation is preserved. It is also important to note the noninvasiveness of OxyBLI in *in vivo* studies; its signals can be collected comprehensively from intact animals because they originate from AkaLumine, a substrate with homogeneous biodistribution^12^. In another approach, O_2_-quenching luminescent dyes, similar to those filled into optical electrodes, can be loaded into the cells of living animals^49^. However, this approach requires high accessibility to objective lenses and involves surgical procedures, such as craniotomy, laparotomy, and thoracotomy, which more or less affect systemic hemodynamics. None of these procedures are required for the basic OxyBLI method.

Second, the genetically-encoded probe of OxyBLI can target any cell type due to advances in gene delivery technology. For example, we monitored OxyBLI signals from neurons specifically, from excitatory vs. inhibitory neurons comparatively (Fig. 2B), or from the brain, limb muscles, and kidneys simultaneously (Fig. 9).

Third, OxyBLI enables rapid, sustained, continuous monitoring of COLs. For example, the use of a conventional EM-CCD camera allowed us to observe OxyBLI signals at 1 Hz and an observation period spanning several hours. We investigated dynamic changes in COLs in a combination of critical situations, including hypoxia, hyperoxia, and anemia, which are typically examined individually. Along with its noninvasive nature, OxyBLI made it possible to test multiple challenges on a single animal over time, revealing some context-dependent aspects of O_2_ regulation (Figs. 5 and 8C). This multifaceted approach will help us better understand the delicate interaction between hypoxia and hyperoxia^50–52^.

Recently, Cai et al. have developed an implantable optoelectronic probe by integrating an O_2_-sensitive phosphorescent film with a light-emitting diode and a photodetector^53^. They implanted the probe in rodents’ brains to continuously monitor PO_2_ in brain tissues. Due to the inherent problem in the O_2_ quenching method, however, the probe was insensitive to high levels of O_2_, thereby limiting its application in monitoring hyperoxia. In fact, the signal rise time of the probe upon a rapid increase in FiO_2_ was determined to be 15.7 sec, too slow for characterizing the overshooting responses that we observed using OxyBLI. In addition, the optoelectronic probe-based approach by Cai et al. appeared to be destructive and unable to correctly detect hyperoxia-induced vasoconstriction responses^53^. In their study, the eventual increment in tissue PO_2_ exceeded 100% at 1.0 FiO_2_. Similar results were obtained in our experiments, in which brain tissue PO_2_ in anesthetized rats was measured using an OxyLite fiber-optic O_2_ probe (Supplementary Fig. 4). The probe implantation likely affected tissue integrity, resulting in loss of normal vascular reactivity to O_2_ and/or contamination of false signals derived from blood vessels.

**Supplementary Table 1.**
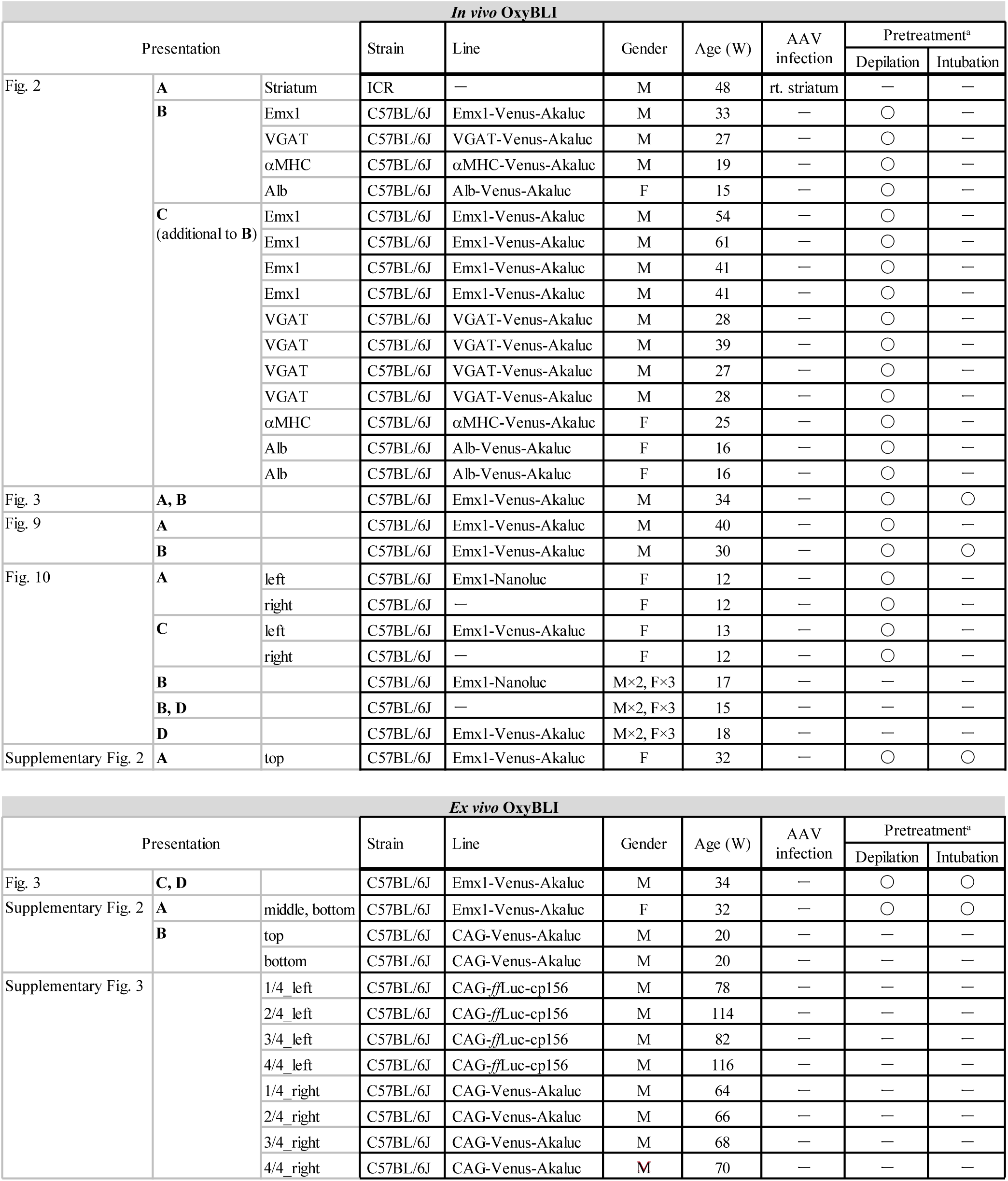
Mice used for OxyBLI (top, *in vivo*; bottom, *ex vivo*) in this study. ^a^Intubation was conducted for artificial ventilation. Figure 3 and Supplementary Figure 2A individually show successive *in vivo*/*ex vivo* BLI experiments.

**Supplementary Table 2.** Rats used for *in vivo* OxyBLI and brain PO_2_ measurement in this study. ^a^Identification numbers were assigned to the rats infected with AAV. ^b^Similar dynamics of the COL signal was observed in the cerebral cortex. ^c^Intubation was conducted for artificial ventilation.

| Presentation |  |  | Strain | Rat ID<sup>a</sup> | Gender | Age (W) | Body Weight (g) | AAV infection | Pretreatment |  |
| --- | --- | --- | --- | --- | --- | --- | --- | --- | --- | --- |
|  |  |  |  |  |  |  |  |  | Depilation | Intubation<sup>c</sup> |
| Fig. 4 | B, C, D |  | SD | #17 | M | 8 | 290 | rt. striatum | ○ | ○ |
| Fig. 5 |  |  | SD | #14 | M | 12 | 386 | rt. striatum | ○ | ○ |
| Fig. 6 | A, B |  | SD | #17 | M | 8 | 290 | rt. striatum | ○ | ○ |
|  | B |  | SD | #14 | M | 12 | 386 | rt. striatum | ○ | ○ |
|  |  |  | SD | #15 | M | 8 | 270 | rt. cortex<sup>b</sup> | ○ | ○ |
|  |  |  | SD | #20 | M | 13 | 470 | rt. striatum | ○ | ○ |
|  |  |  | SD | #48 | M | 15 | 625 | rt. striatum | ○ | ○ |
|  |  |  | SD | #53 | M | 12 | 430 | rt. striatum | ○ | ○ |
| Fig. 7 | A, B |  | SD | #41 | M | 11 | 422 | rt. striatum | ○ | ○ |
| Fig. 8 | A |  | SD | #20 | M | 12 | 470 | rt. striatum | ○ | ○ |
|  | B |  | SD | #35 | M | 12 | 400 | rt. striatum | ○ | ○ |
|  | C |  | SD | #14 | M | 12 | 386 | rt. striatum | ○ | ○ |
| Supplementary Fig. 4 | B |  | SD | — | M | 13 |  | — | — | — |
|  | C |  | SD | — | M | 17 |  | — | — | — |

**Supplementary Table 3.**
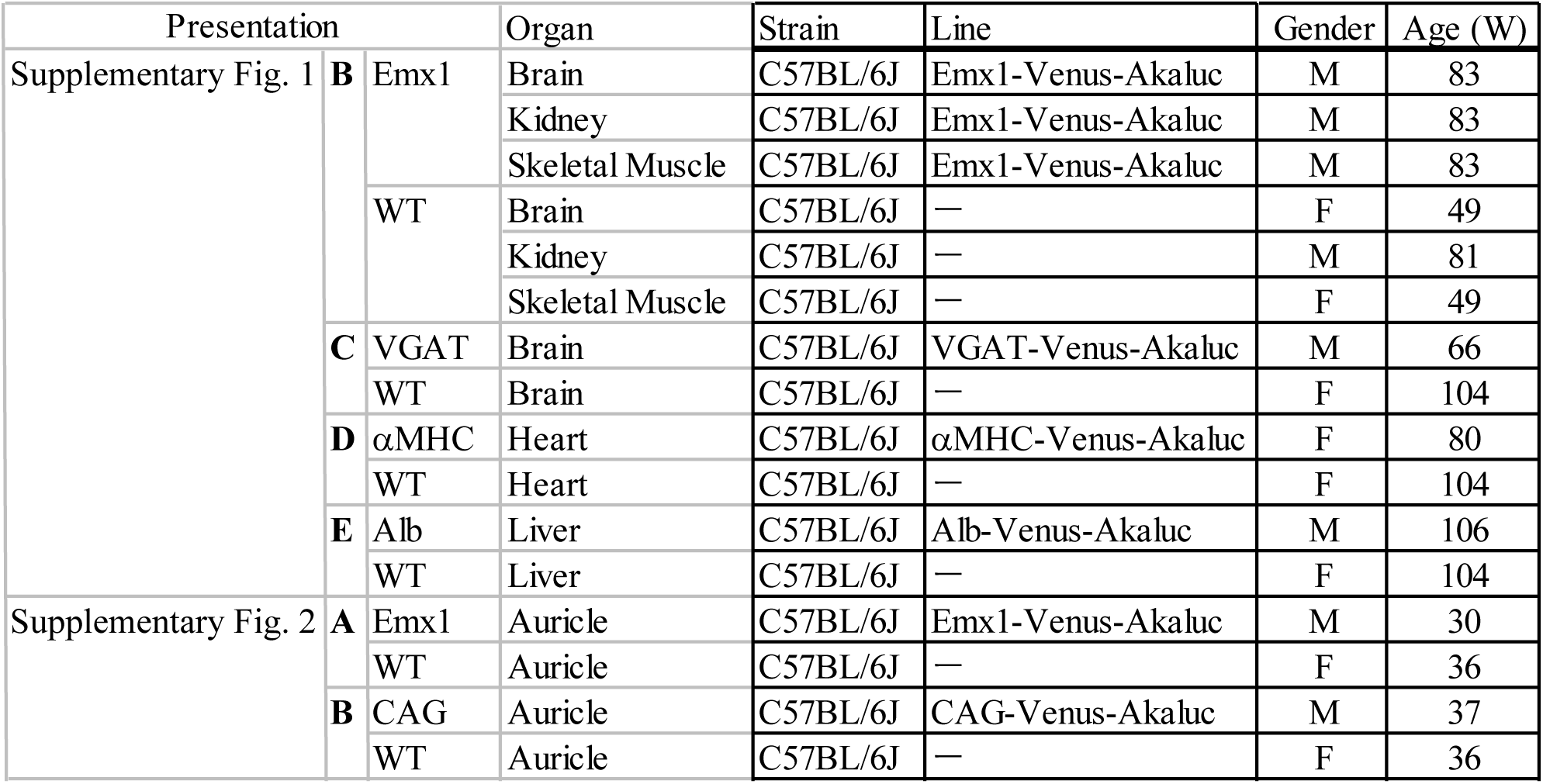
Mice used for immunohistochemistry in this study.

**Supplementary Figure 1.**
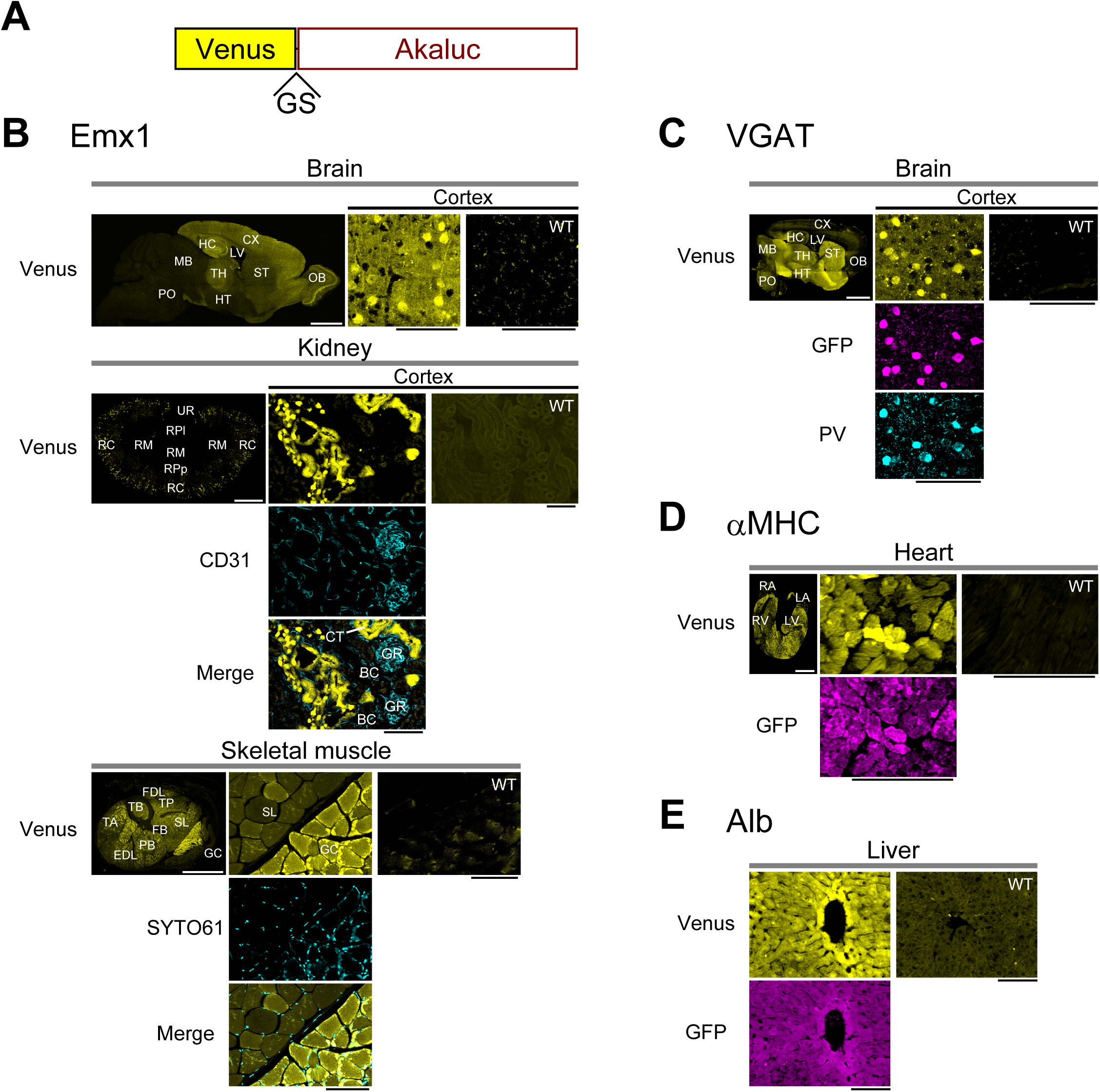
Spatial patterns of Akaluc expression visualized by Venus fluorescence. **A,** Domain structure of Venus-Akaluc fusion. GS: glycine and serine. **B–E,** Tissue sections from Emx1-Venus-Akaluc (**B**), VGAT-Venus-Akaluc (**C**), αMHC-Venus-Akaluc (**D**), and Alb-Venus-Akaluc (**E**) mice were imaged for Venus fluorescence. In parallel, similar sections from wild-type (WT) mice were imaged in the same way. White scale bars: 2 mm. Black scale bars: 100 μm. **B,** Distinct Venus fluorescencewas observed in all the expected excitatory neurons, all the skeletal muscle fibers in the antebrachial part, and a fraction of proximal tubeles in the renal cortex. Kidney and skeletal muscle sections were immunostained for CD31 and counterstained for nuclei (SYTO61), respectively. **C,** Brain sections were immunostained for parvalbumin (PV), a marker of inhibitory interneurons. **D,** Heart sections. **E,** Liver sections. **C–E,** Sections were immunostained for Venus by using an anti-GFP antibody. Abbreviations (brain) in (**B**) and (**C**): CX: cerebral cortex; HC: hippocampus; HT: hypothalamus; LV: lateral ventricle; MB: midbrain; OB: olfactory bulb; PO: pons; ST: striatum; TH: thalamus. Abbreviations (kidney) in (**B**): BC: Bowman’s capsule; CT: collecting tubule; GR: glomerulus; RC: renal cortex; RM: renal medulla; RPl: renal pelvis; RPp: renal papilla; UR: ureter. Abbreviations (skeletal muscle) in (**B**): EDL: extensor digitorum longus; FB: fibularis; FDL: flexor digitorum longus; GC: gastrocnemius; PB: peroneus brevis; SL: soleus; TA: tibialis anterior; TB: tibia; TP: tibialis posterior. Abbreviations (heart) in (**D**): LA: left atrium; LV: left ventricle; RA: right atrium; RV: right ventricle.

**Supplementary Figure 2.**
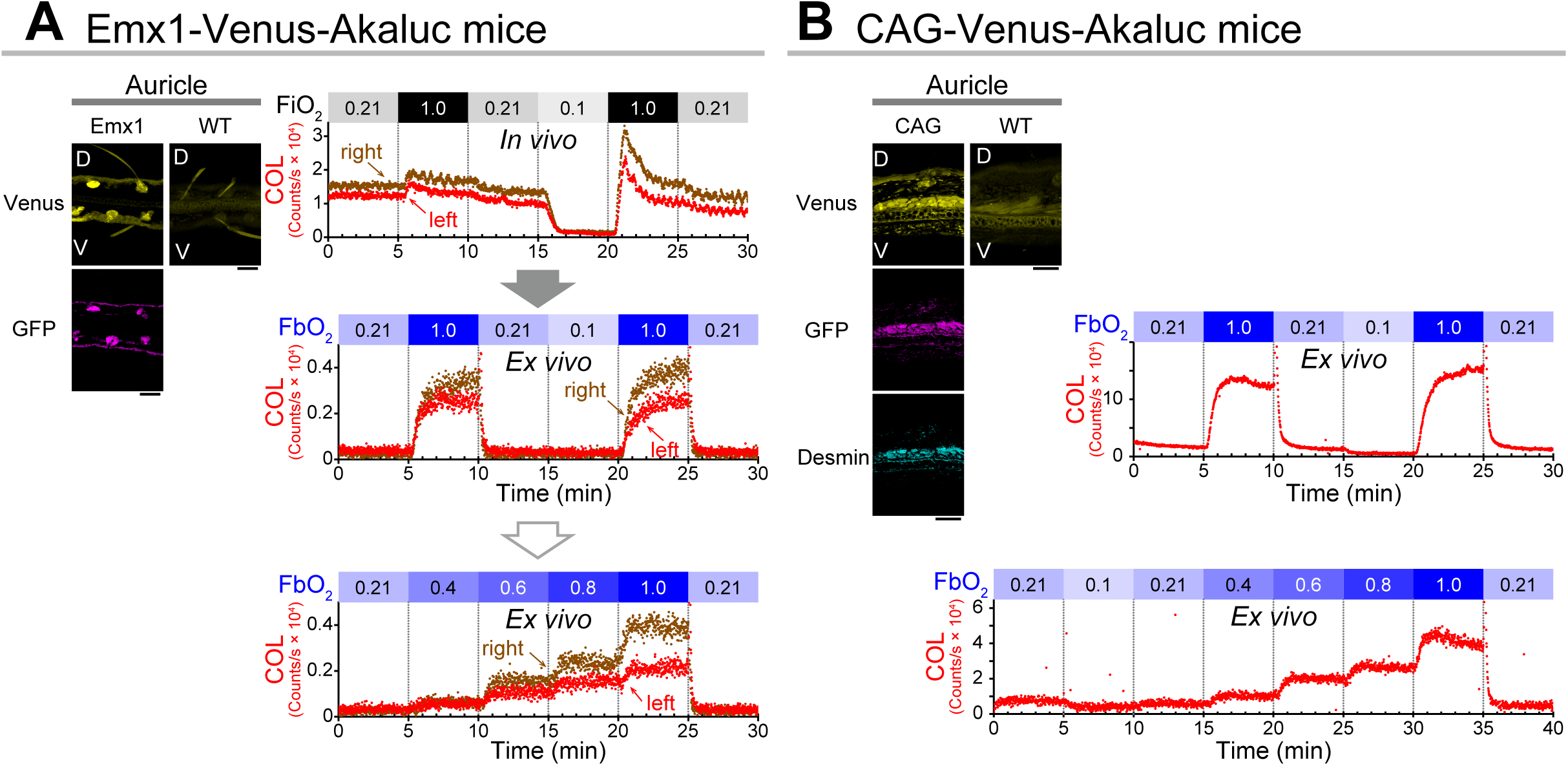
Cellular O_2_ levels (COLs) in detached mouse auricles. **A,** Auricles were prepared from Emx1-Venus-Akaluc mice for OxyBLI. *left,* Fluorescence (FL) images of auricular transverse sections from Emx1-Venus-Akaluc (Emx1) and wild-type (WT) mice. The Emx1 section was reacted with 1^st^ Ab (rabbit Ab to GFP) and 2^nd^ Ab (goat Ab to rabbit IgG conjugated to DyLight 550). No reactions were conducted on WT sections. Representative confocal images showing Venus (top) and DyLight 550 (bottom) fluorescence. Autofluorescence of hair shafts is evident in the WT image. D, dorsal side. V, ventral side. Scale bars: 0.1 mm. *right,* Successive *in vivo*/*ex vivo* imaging. A gray arrow indicates the transfer of BLI from *in vivo* to *ex vivo*. After detachment, the left and right auricles were placed on clay, as done in the experiments depicted in Figure 3, but neither the position nor the lens of the camera was altered. As a result, the COL data were relatively noisy owing to weak signals compared with those seen in Figure 3D. Two FbO_2_ challenges (the regular protocol and then the stepwise increases) were conducted sequentially (an open arrow). The fraction of inspired O_2_ (FiO_2_) and the fraction of blown O_2_ (FbO_2_) are indicated by shades of gray and blue, respectively. **B,** Auricles were prepared from CAG-Venus-Akaluc mice for OxyBLI. *left,* Fluorescence (FL) images of auricular transverse sections from CAG-Venus-Akaluc (CAG) and wild-type (WT) mice. The CAG section was reacted with rabbit Ab to GFP and mouse Ab to Desmin, and then visualized with goat Ab to rabbit IgG conjugated to DyLight 550 and donkey Ab to mouse IgG conjugated to Alexa Fluor 647, respectively. The immunofluorescence for Desmin highlights the middle muscular layer. D, dorsal side. V, ventral side. Scale bars: 0.1 mm. *right,* Two *ex vivo* experimental data from different mice. Time courses of COL in the auricle during the regular FbO_2_ protocol (top) and an FbO_2_ challenge that included a 5-min hypoxia at 0.1 and stepwise increases from 0.21 to 1.0 (bottom). The fraction of blown O_2_ (FbO_2_) is indicated by shades of blue. Given the localization of Venus-Akaluc in the middle muscular layer of the auricle, under the control of the CAG promoter^54^ it is suggested that O_2_ diffuses into the center of this auricular part, which has a thickness of a few hundred micrometers.

**Supplementary Figure 3.**
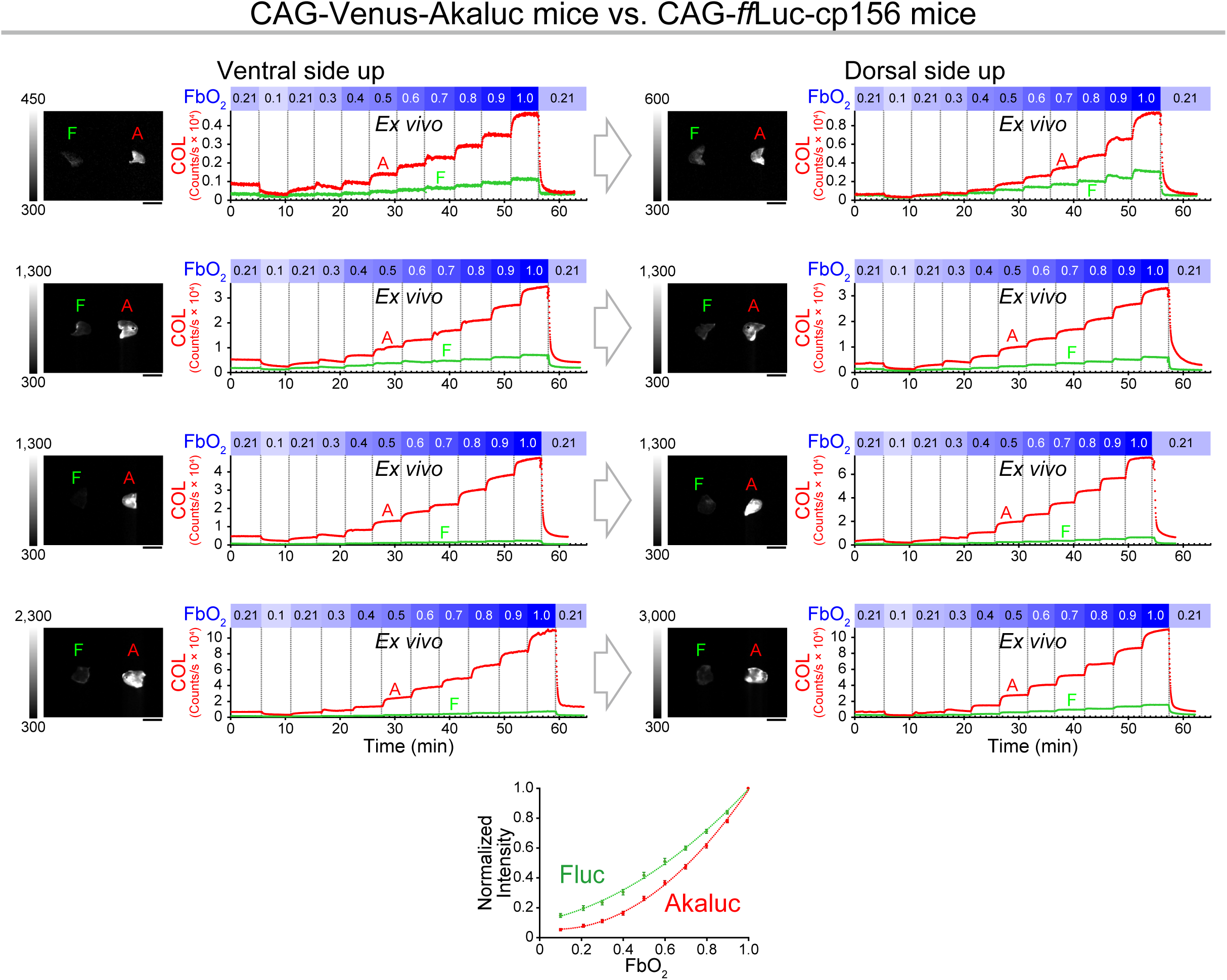
Dose (O_2_)-response (emission) relationships of Fluc vs. Akaluc. Comparative *ex vivo* experiments using CAG-*ff*Luc-cp156 and CAG-Venus-Akaluc mice after ip administration of D-luciferin and AkaLumine-HCl, respectively. In each setting, a pair of auricles detached from the two lines was subjected to two consecutive gas exposures (an open arrow) with stepwise increases in FbO_2_. The gases were blown onto the ventral and dorsal sides of the auricles in the first (left) and second (right) exposures, respectively. Four pairs were used for the execution of 8 independent experiments. Images: *left*, Auricles from CAG-*ff*Luc-cp156 mice (F); *right*, Auricles from CAG-Venus-Akaluc mice (A). Scale bars: 10 mm. Temporal profiles of auricular COL are shown in green (Fluc) and red (Akaluc). The fraction of blown O_2_ (FbO_2_) is graded by shades of blue. The O_2_ dependencies of Fluc (green) and Akaluc (red) are summarized at the bottom. Normalized to the intensities at 1.0 FbO_2_. Fluc data were fitted to the equation y = 0.601 x^2^ + 0.2824 x + 0.1091. Akaluc data were fitted to the equation y = 1.088 x^2^ – 0.166 x + 0.0623. Data points are shown as mean ± s.e.m. (*n* = 8).

**Supplementary Figure 4.**
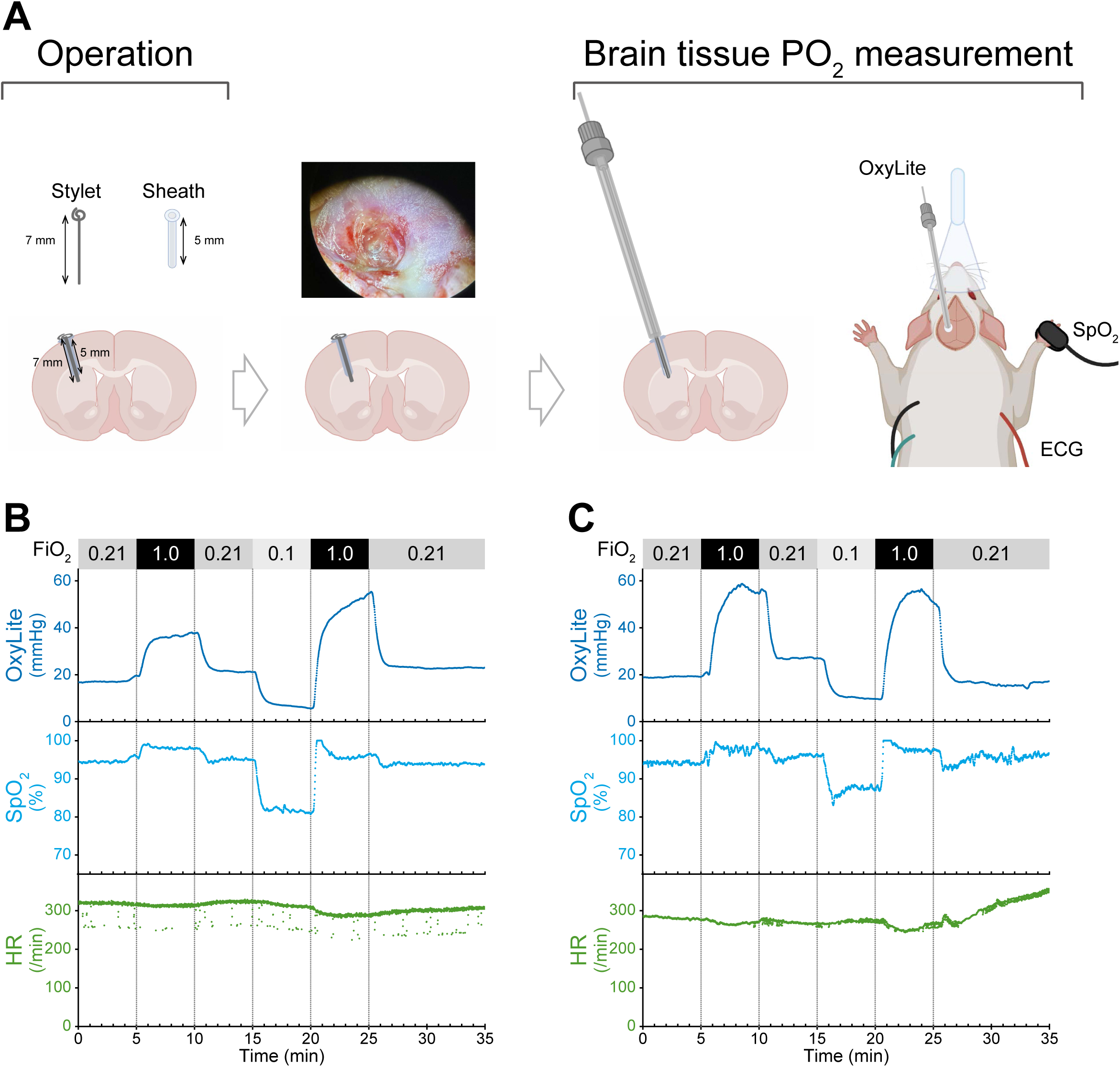
An optoelectronic probe-based approach for measuring rat brain tissue PO_2_ under hypoxic and hyperoxic conditions. **A,** Experimental design. *left,* Surgery was done to implant a plastic sheath into the left brain of a wild-type rat. With this sheath, a metal stylet was inserted with its tip located 7 mm from the brain surface. *middle,* Several days post-operation, it was possible to confirm by appearance that the inflammation had subsided. *right,* the OxyLite fiber-optic O_2_ probe was substituted for the stylet to measure interstitial PO_2_ in the striatum. The rat was equipped with an ECG and a pulse oximeter for monitoring HR and SpO_2_, respectively. **B,C,** Time courses of brain tissue PO_2_ (OxyLite), SpO_2_, and HR during the regular FiO_2_ protocol. Two datasets using different rats are depicted, which were obtained 6 (**B**) and 23 (**C**) days after the operation. Hyperoxic breathing (1.0 FiO_2_) increased the OxyLite signals to a maximum, approximately 100–200% above the baseline. The fraction of inspired O_2_ (FiO_2_) is indicated by shades of gray.

**Supplementary Video 1. Visualization of COLs in spontaneously breathing mice under hypoxic and hyperoxic conditions.**

The movies of the following mice are played in sequential order: a mouse with striatal expression of Venus-Akaluc (see Fig. 2A); an Emx1-Venus-Akaluc mouse (see Fig. 2B); a VGAT-Venus-Akaluc mouse (see Fig. 2B); an αMHC-Venus-Akaluc mouse (see Fig. 2B); an Alb-Venus-Akaluc mouse (see Fig. 2B).

The COL time courses are shown at the bottom. For the Emx1-Venus-Akaluc mouse, the ROI was localized to the head to monitor the brain COL. This video (2.6 MB) has been generated via considerable compression of the original large-volume video data (0.40 GB). Compression was made using FFmpeg.

**Supplementary Video 2. Side-by-side comparison of *in vivo* and *ex vivo* COL visualization during the regular FiO_2_ protocol.**

The movies of the whole body (left) and detached auricle (right) correspond to Figure 3B and Figure 3D, respectively, and are played simultaneously. This video (1.9 MB) has been generated via considerable compression of the original large-volume video data (0.35 GB). Compression was made using FFmpeg.

**Supplementary Video 3. Visualization of COL in the rat striatum during the regular FiO_2_ protocol.**

See Figure 4D. The COL time course is shown at the bottom. This video (9.5 MB) has been generated via considerable compression of the original large-volume video data (1.62 GB). Compression was made using FFmpeg.

